# Genetic variation in recalcitrant repetitive regions of the *Drosophila melanogaster* genome

**DOI:** 10.1101/2024.06.11.598575

**Authors:** Harsh G. Shukla, Mahul Chakraborty, J.J. Emerson

## Abstract

Many essential functions of organisms are encoded in highly repetitive genomic regions, including histones involved in DNA packaging, centromeres that are core components of chromosome segregation, ribosomal RNA comprising the protein translation machinery, telomeres that ensure chromosome integrity, piRNA clusters encoding host defenses against selfish elements, and virtually the entire Y chromosome. These regions, formed by highly similar tandem arrays, pose significant challenges for experimental and informatic study, impeding sequence-level descriptions essential for understanding genetic variation. Here, we report the assembly and variation analysis of such repetitive regions in *Drosophila melanogaster*, offering significant improvements to the existing community reference assembly. Our work successfully recovers previously elusive segments, including complete reconstructions of the histone locus and the pericentric heterochromatin of the X chromosome, spanning the *Stellate* locus to the distal flank of the rDNA cluster. To infer structural changes in these regions where alignments are often not practicable, we introduce landmark anchors based on unique variants that are putatively orthologous. These regions display considerable structural variation between different *D. melanogaster* strains, exhibiting differences in copy number and organization of homologous repeat units between haplotypes. In the histone cluster, although we observe minimal genetic exchange indicative of crossing over, the variation patterns suggest mechanisms such as unequal sister chromatid exchange. We also examine the prevalence and scale of concerted evolution in the histone and Stellate clusters and discuss the mechanisms underlying these observed patterns.

## Introduction

Tandem repetitive sequences are major components of many eukaryotic genomes (Richard et al. 2008) and play vital roles in both biology (Gemayel et al. 2010; Hannan 2012) and disease (Hannan 2018). Loci encoding components of important cellular processes are sometimes present in multiple copies and are often clustered in tandem, including examples like histone genes (Doenecke 2014), rDNA arrays (Hall et al. 2022), and immunoglobulin genes (Watson and Breden 2012). Tandem repeats are prone to rapid copy number changes mediated through mechanisms such as unequal crossing over, replication slippage, gene conversion, and intra-chromatid homologous recombination (Smith 1976; Charlesworth et al. 1994; Johnson and Jasin 2000; Cohen et al. 2003). They also evolve rapidly, as evidenced by the rapid turnover of satellites both in abundance (copy number) as well as specific repeat types between species (Wei et al. 2018; Cechova et al. 2019). Elucidating the mutational mechanisms and evolutionary forces acting on the tandem arrays requires reliably discovering and distinguishing alleles within and between individuals (the “fundamental datum” in (Hubby and Lewontin 1966; Lewontin and Hubby 1966; Charlesworth et al. 2016)). However, most contemporary reference genome assemblies, including those resulting from consortium-led efforts, contain gaps in such repetitive regions (Altemose et al. 2014; Hoskins et al. 2015; Nurk et al. 2022). This limitation constrains routine comparisons between tandem repeat arrays necessary to elucidate the structure, function, and evolution of such arrays.

The rapid development of long-read sequencing technologies (Marx 2023) and informatics approaches (Chin et al. 2013; Berlin et al. 2015; Chin et al. 2016; Koren et al. 2017) led to a proliferation of genome assemblies that approached the quality of consortium-led reference sequencing projects. In *Drosophila*, these advances led to enormous improvements in the ability of single research groups to rapidly and inexpensively construct high-quality genome assemblies (Chakraborty et al. 2016; Chakraborty et al. 2018; Mahajan et al. 2018; Miller et al. 2018; Solares et al. 2018; Chakraborty et al. 2019; Hill et al. 2019; Chakraborty et al. 2021; Kim et al. 2021; Liao et al. 2021). But, owing to its inherently noisy nature, some repetitive parts of the genome were still recalcitrant to assembly approaches (Nurk et al. 2020; Rhie et al. 2021). While sophisticated application of long read data to assembly problems has unarguably led to several examples of advancing the state of the art in previously recalcitrant regions of the genomes (Khost et al. 2017; Philipp Bongartz and Schloissnig 2019; Chang and Larracuente 2019), these approaches were not scalable to routine application by typical research groups, requiring extensive novel work on assembly informatics before downstream analysis was possible. Indeed, despite the promise of long read approaches, cytogenetic approaches remain indispensable to studying genetic variation in such regions (Courret and Larracuente 2023).

Consequently, studies of variation in many parts of the genome harboring tandem repeat arrays remain challenging. It has long been recognized that copies of a gene in a particular species may be more closely related than orthologs in related species, leading to species-specific variants in gene families that predate species divergence (Hood et al. 1975; Liao 2008). This homogenization of repeat units, termed concerted evolution (Liao 2008), is mediated through DNA recombination, repair, and replication mechanisms such as unequal crossing over and gene conversion (Liao 1999). However, the extent of homogenization and variation in observed copy numbers may vary across tandem arrays and may be shaped by various factors (such as natural selection, drift, mutation, drive, age, size, and relative rates of different kinds of recombination) acting on the array (Dover et al. 1993; Elder and Turner 1995; Lower et al. 2018). Untangling patterns of concerted evolution on tandem arrays experiencing different forces requires a comprehensive map of genetic variation spanning such clusters.

The cluster of highly conserved Histone genes on chromosome arm 2L and the species-specific X-linked cluster of *Stellate* (*Ste*) genes are examples of two recalcitrant tandem arrays in *D. melanogaster* shaped by different evolutionary forces. The histone locus has a tandem organization with the quintet of 5 histone genes (H1, H2B, H2A, H4, H3, from distal to proximal) arrayed in tandem for ∼100-110 times (Lifton 1978; Strausbaugh and Weinberg 1982). Each unit is ∼5 kb in length, and this tandem organization is present in distantly related *Drosophila* species, indicating old evolutionary origins (Kakita et al. 2003). In contrast, Stellate is a *D. melanogaster*-specific, evolutionary young tandem cluster thought to have arisen through the intragenomic conflict between X & Y chromosomes (Malone et al. 2015; Martí and Larracuente 2023). It is present at two distinct locations on the X chromosome: one in euchromatin and the other in heterochromatin, and each unit is 1.26 kb and 1.15 kb in size, respectively (Palumbo et al. 1994; Adashev et al. 2020). In wild-type male files, the *Stellate* locus is normally suppressed by piRNAs originating from the complementary *Su(Ste)* loci on the Y chromosome. If left unsuppressed, *Stellate* RNA produces star/needle-like crystal protein aggregates in primary spermatocytes, leading to meiotic abnormalities and fertility defects (Adashev et al. 2020). Studying genetic variation within these two tandem arrays can help us understand the general and unique properties of the mutational mechanism and concerted evolution driving the evolution of these complex regions of the genome. However, we lack a detailed map of genetic variation within tandem arrays like Histone or Stellate clusters, limiting inferences of molecular mechanisms or patterns of evolution in such clusters.

Here, we present analyses of genome structure for three strains of *D. melanogaster* that address such challenging regions of the *D. melanogaster* genome, including the Histone and Stellate clusters. To develop a scalable approach for studying genetic variation within tandem arrays, we applied widely used software to easily collectable highly accurate long-read data (Wenger et al. 2019) to assemble the recalcitrant repetitive regions of the genome and developed a new framework to reveal the detailed pattern of structural variation in repeat clusters. Using a comprehensive map of variation within the Histone and Stellate clusters, we explore patterns of variation that empower us to comment on mutational processes shaping two functionally important tandem arrays in the *Drosophila* genome.

## Results

### Genome Assemblies

The strains selected include the *Drosophila* community’s genome reference strain (iso-1) (Adams et al. 2000) and two strains from the *Drosophila* Synthetic Population Resource (A3 and A4) (King et al. 2012), all of which are near isogenic. These were sequenced with Pacbio HiFi (Wenger et al. 2019) sequencing chemistry. The assemblies are very contiguous and complete, exhibiting contig N50s (L50s) of 22.96 Mb (4), 21.53 Mb (4), and 21.50 Mb (4) for the iso-1, A4, and A3 assemblies, respectively, as well as single-copy BUSCO scores of 99.42%, 99.27%, and 99.42%. The assemblies exhibit very few base-level errors, exhibiting phred (QV) scores of 49.9, 56.1, and 55.3 with estimates of heterozygosity for these isogenic strains of 0.076%, 0.071%, and 0.081% for iso-1, A4, and A3, respectively. The chromosome arms are mostly represented by single contigs spanning the entire euchromatin (Fig 1A). Gaps appear in highly repetitive heterochromatic regions of the genome (Fig 1A; red boxes occurring in pericentromeric regions of chromosomes 2 and 3). The median read length (15-20 kb) of HiFi sequences is not sufficient to span many satellites and other repetitive sequences (Porubsky et al. 2023) enriched in the pericentromeric and centromeric heterochromatin (Jagannathan et al. 2017; Shatskikh et al. 2020). The iso-1 HiFi assembly was scaffolded using both HiC data and reference-assisted scaffolding (Supp Fig 4). In autosomes (Muller elements B-F), ∼4.17 Mb of new sequence was added relative to Release 6, most of it (∼3.29 Mb) in heterochromatic regions (Fig 1C). For the X chromosome (Muller element A), our assembly also introduced ∼3.9 Mb of new sequence, including ∼2.76 Mb of heterochromatin sequence adjacent to the rDNA array. For every major repeat category across all chromosomes except the dot, the HiFi assembly of iso-1 recovered more than previous assemblies (Fig 1B). We also identified a total of 15.7, 16.48, and 17.03 Mb of contigs putatively belonging to the Y chromosome in iso-1, A4, and A3, respectively. The iso1 Y assembly corresponds well in structure and repeat content to a recently reported assembly (Chang and Larracuente 2019) (Supp Fig 5). The assemblies of A3 and A4 were scaffolded using comparative scaffolding with our iso-1 HiFi scaffolded assembly as the reference. The assemblies exhibited scaffold N50s (L50s) of 27.44 Mb (3), 27.72 Mb (3), and 28.05 (3) for iso-1, A4, and A3, respectively. In total, the chromosome scaffold for all three strains spans ∼142 Mb. These assemblies are superior to existing high-quality reference-grade assemblies like iso-1 Release 6 (Fig 1D). The statistics of the data and assembly generated can be found in Supp. Table 1-5.

**Fig. 1:**
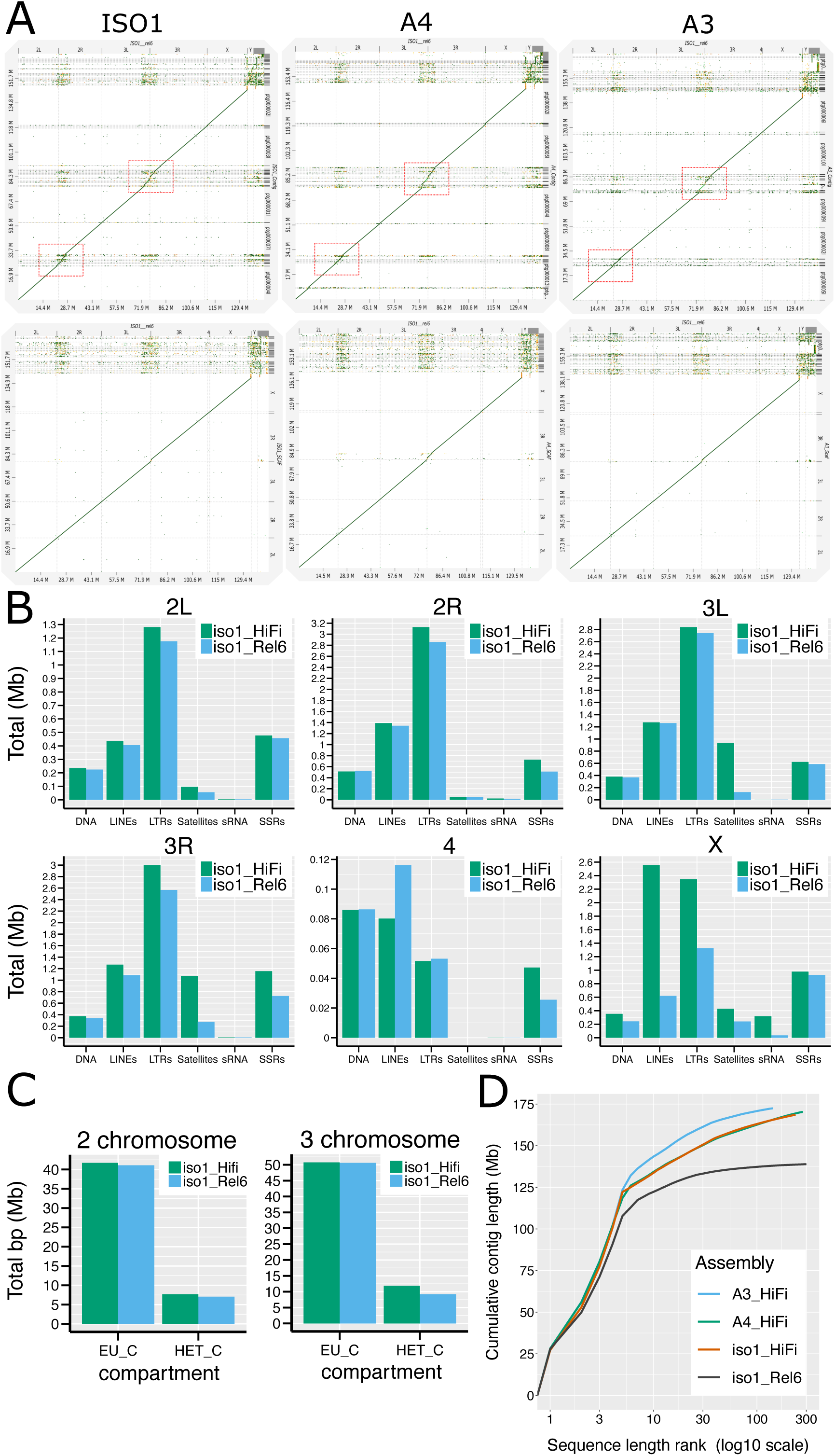
Comparison of assemblies here to that of Release 6. **A.** D-Genies dotplots of the Release 6 reference genome assembly scaffolds for Drosophila melanogaster (x-axis) versus our contig-(top) and scaffold-level (bottom) assemblies of iso-1, A4 and A3. **B.** Repeat content comparison of iso-1 Release 6 assembly versus our iso-1 HiFi assembly for each Muller element (i.e., autosomal chromosome arms and the X chromosome). **C.** Comparison of total sequence assigned to euchromatic and heterochromatic compartments in iso-1 Rel6 versus iso-1 HiFi. **D.** Cumulative length plot for assembly contigs of strains used in this study and iso-1 Release 6. The x-axis is on a log_10_ scale.

### Newly assembled X-linked heterochromatic sequence

The heterochromatic portions of the genome have been subdivided into their own bands based on cytological features (Kaufman 2017). The heterochromatin of the X chromosome spans bands *h26-h34*, out of which only a small part of *h26* is represented in the chromosome-scale scaffolds belonging to iso-1 assembly on FlyBase. Our iso-1 assembly recovers a single contig spanning the entire X euchromatin and heterochromatic bands *h26-h28* and 454 kb of *h29*, the band containing the rDNA gene cluster. Alignments of the X chromosome between HiFi assemblies from iso-1, A4, and A3 to Release 6 (Fig 2A) show that the newly assembled segments *h26-h28* comprise arrays of segmental duplications with varying copy numbers in the 3 strains. The entire region (∼98.5%) is repetitive, with two repeat elements – LINE R1/R2 – dominating the landscape (Fig 2B). Fig 2C compares the whole region across the 3 assemblies. The iso-1 and A3 strains exhibit similar haplotype structures, except A3 harbors one additional large multi-kb segment. The A4 haplotype varies considerably from the other two, with only part of the proximal and distal ends of the array aligning. The intra-strain self-dot plots illustrate this, with iso-1 and A3 (Supp Fig 6) exhibiting similar patterns while A4 is quite different. To verify the integrity of the newly assembled region, we mapped HiFi data back to the assembly and inspected the depth in this region. Dramatic variation in depth plots may indicate errors in the assemblies, such as putative misassemblies, collapsed or expanded duplications, etc. The depth plot (Supp Fig 7) for iso-1 indicates that a region of ∼250 kb immediately distal to the rDNA exhibits twice the coverage, indicating possible collapse in our assembly. The plots for A4 and A3 (Supp Fig 7) show relatively uniform coverage, indicating no large-scale mis-assemblies.

**Fig. 2:**
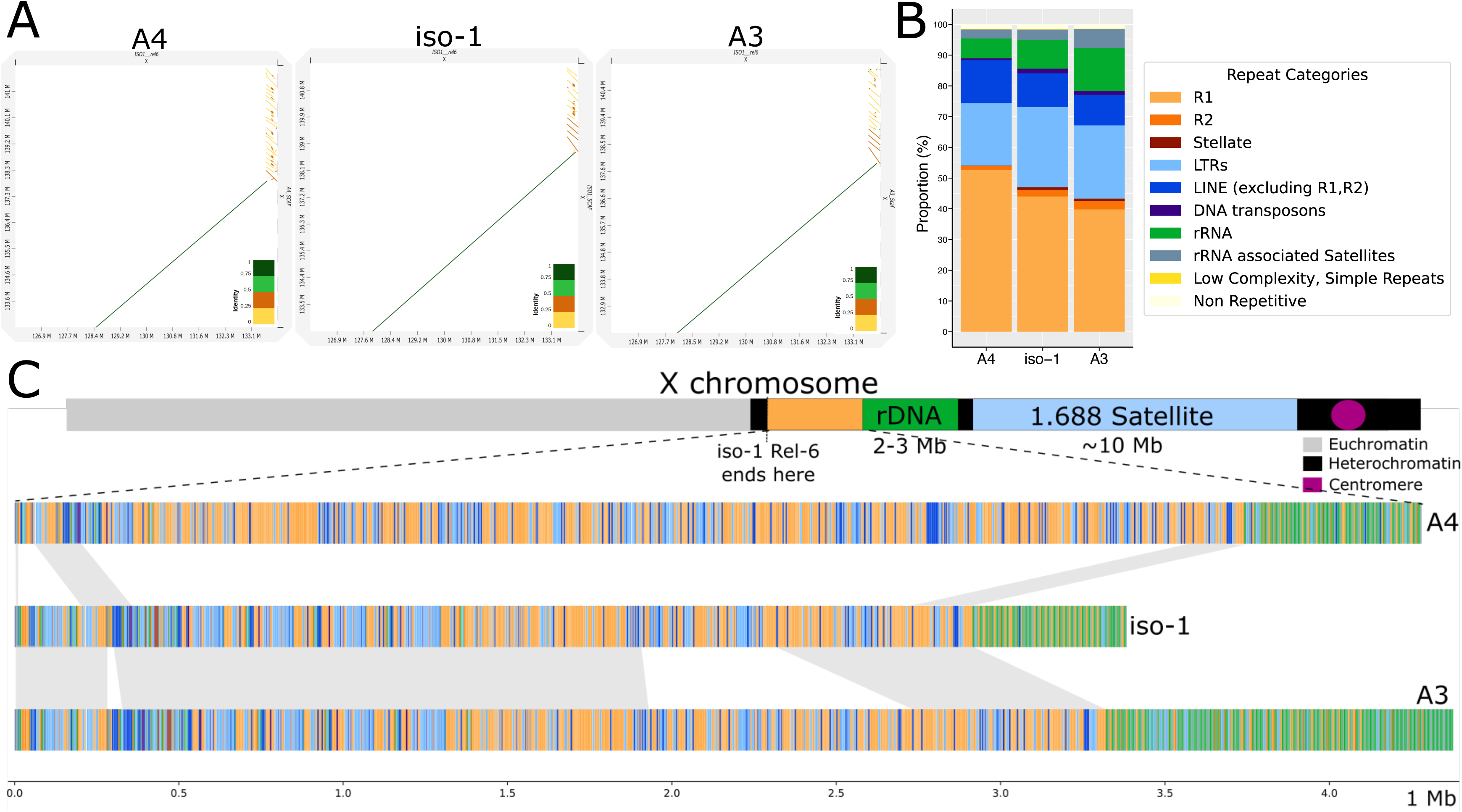
Characterization of newly assembled X-linked heterochromatic sequence. **A.** D-Genies dotplots of our HiFi assemblies (y-axis) versus the Release 6 assembly of iso-1 (x-axis) covering the proximal end of X chromosome scaffold for both assemblies. **B.** Repeat catalog of the newly assembled proximal regions on the X chromosome for different strains. **C.** Map of sequence from the newly assembled proximal end of the contigs assembled here. Top: Schematic map of X chromosome structure. Bottom: structure of the region in the HiFi assemblies of the X chromosome for the haplotypes in A4, iso-1, and A3. The haplotypes are painted with colors representing different repeat categories (following panel B).

### Structural variation in the histone cluster

#### Assembly of *D. melanogaster* histone cluster for iso-1, A4, and A3

The histone locus in *Drosophila melanogaster* is present at the 39DE region of chromosome 2L, bordering the pericentromeric heterochromatin. The highly homogenous nature of this locus has impeded its resolution to the base pair level. The current reference iso-1 Rel6 has only ∼69 kb (12 units) assembled on the distal flank and ∼56 kb (11 units) assembled on the proximal flank. Our default assembly model of the histone locus in iso-1 spans the entire locus without gaps. The array is ∼577kb, comprising a total of 111 units. This is consistent with expectations from the literature. We mapped the raw HiFi read data back to the assembly to quantify the read coverage across the locus and validate this result. In a well-assembled region, the read coverage should be uniform, lacking sudden and dramatic changes in coverage in a short span. The sudden drop and rise in coverage and doubled coverage (at ∼200kb and ∼50kb) suggest the presence of a misassembly and a small collapse at the beginning of the cluster, respectively (Supp Fig 8). To address these features, we employed a targeted assembly approach where we identified reads mapping to the histone locus. We then assembled them with parameters more appropriate for repeats (See Methods). The targeted assembly exhibited a locus size of ∼587kb and 113 copies. The coverage plot of the targeted assembly is comparatively uniform, with no sudden rise and drop in coverage, suggesting that the misassembly is no longer present (Supp Fig 8). It also no longer possesses the doubled coverage indicative of a collapsed duplication. The putative collapsed tandem duplication spanned 2 units (units 7-8 and 9-10 in the targeted assembly), increasing the total unit copy number by two in the targeted assembly. The putative misassembly affected unit 39 (unit 37 in the original assembly). Other than these differences, the structure of both assemblies remains the same between the default assembly and the targeted assembly. The assembly graph around the histone locus in the original assembly had only 3 unitigs (Supp Fig 9A). They overlap with each other at the same junction, with smaller unit 842l having an overhang. Thus, there was only one possible path through it. This junction is near the collapsed duplication, suggesting a cause for the collapse. Though the unitig graph of the targeted assembly had more unitigs, the path through it was relatively clear. (Supp Fig 9B)

We compared our assembly of iso-1 to that of Bongartz et al. (P. Bongartz and Schloissnig 2019), who employed machine learning to assemble the histone cluster using published long-read data (Kim et al. 2014). The structure of this assembly (henceforth referred to as iso-1-BS19) was mostly consistent with ours, with units 2-107 of iso-1-BS19 corresponding to units 6-111 of ours. All discrepancies between the assemblies were located at the edges of the cluster (Supp Fig 10). Units 1-4 of our assembly were missing in iso-1-BS19. The iso-1-BS19 starts with a partially assembled unit containing *LTR/roo* corresponding to the 5th copy in our assembly. On the other side of the cluster, the *DNA/Pogo*-containing unit is the final unit (113) in our HiFi assembly, whereas it is the penultimate copy (unit 108) in the iso-1-BS19 version. Instead of the final copy (corresponding to unit 113 in our assembly), iso-1-BS19 has a partially complete unit (∼3.3kb). By aligning the two assemblies across ∼105 units (corresponding to units 6-111 in our assembly and alignment block length of ∼540 kbp), we discovered only two SNPs and two 1-bp indels associated with homopolymer tracts of length 8 and 13.

The default assembly for A4 constructed the histone cluster into 119 copies. The read depth plot shows a drop in coverage (about half) for around ∼40-50 kb towards the end of the cluster (Supp Fig 11). This indicates that a region present once in the genome is represented twice in the assembly. The unitig graph (Supp Fig 12A) in that region shows two unitigs (963l and 738l) that overlap on both ends. It appears that *hifiasm* visited the 963l node twice, leading to the duplication. The targeted assembly of the histone locus has 111 copies, totaling ∼571 kb. The read depth plot shows relatively uniform coverage and no apparent misassembly or collapse/expansion (Supp Fig 11). The previous drop in coverage also disappears. The targeted unitig graph has a slightly complex region in the beginning, but *hifiasm* resolves it (Supp Fig 12B).

The default assembly for A3 constructed the histone cluster into 114 copies. Depth plots identified a putative misassembly at ∼200kb immediately followed - by several large collapsed regions (Supp Fig 13). The default unitig graph around the histone locus is more complex than either iso-1 or A4 (Supp Fig 14A). Moreover, a targeted assembly with modified parameters failed to assemble the cluster, exhibiting breaks, suggesting that the structure of A3’s histone cluster possessed features more recalcitrant to assembly than either iso-1 or A4, like large and homogeneous duplications. This also hinted that the histone unit copy number was significantly higher than the typical expectation of ∼110. To validate our findings, we sequenced PCR-free Illumina libraries derived from the 3 strains (the same ones in our lab that were used to generate HiFi data). We adapted the idea developed by (McKay et al. 2015) for estimating copy numbers. Briefly, we masked the entire histone locus in the iso-1 hifi assembly, introduced only a single copy of the histone unit as a separate contig, and mapped Illumina data to it. The relative ratio of the depth of the histone contig (1 unit) vs the depth of any single copy region should approximate the total number of histone units present in the genome. Such an estimate would be a lower bound since only one unit is present, and any variation between units could lead to reduced mapping. We measured the ratio of the median depth of the histone contig to the median depth of the 2L chromosome. Both iso-1 and A4 showed results consistent with, but slightly lower than, observed in the assemblies (102 for iso-1 and 102 for A4). However, the A3 coverage showed an elevated copy number of 197, almost double the typical copy number. Because the standard targeted assembly approach for A3 did not yield a contiguous assembly, we subsampled the longest 38Mb of reads, which would be 40X if the locus were 950kb (∼190 units) in length. These reads were then used to construct a final targeted assembly based on longer reads, which led to three contigs spanning ∼134 units, which is still shy of our conservative prediction of 197. As a result of this inability to reconstruct the entire histone cluster in A3 without gaps, downstream analyses focus on only the well-resolved parts of the locus (Supp Fig 14B). The unitig graph from the longest 40X data shows a very complex region in the middle exhibiting possibly circular paths (Supp Fig 14B). This likely corresponds to large duplications indicated by the read depth plots. The correct path through it cannot be unambiguously determined. For our final assembly, we considered the well-resolved parts on the left (∼193 kb) and right (∼307kb) flanks and joined them with a gap containing Ns. The total size of our gap-containing model of the locus was ∼500 kb and had 97 units (36 on the left flank and 61 on the right).

#### Inferring histone cluster copy numbers in the global diversity lines

To estimate the copy number in the histone cluster in stocks when no high-quality assembly exists, we used the depth of Illumina data. We focused on a reference collection of 85 *Drosophila* stocks representing genetic diversity across the globe (i.e., the Global Diversity Lines or GDL) (Grenier et al. 2015). The mean and median histone copy numbers in GDL were 101.27 and 104, respectively. The lowest copy number we observed was 29, and the highest was 146. The standard deviation was 20.56. Supp Fig 15 shows the distribution of inferred copy numbers in GDL.

We took advantage of replication and deep sequencing in the reference collection to estimate the error in copy number estimated from our approach. One stock (ZW155-ST/ST) was sequenced twice, once to 12.5X (the average depth for all samples) and once to a depth of ∼100X. Our pipeline estimated 81 copies for the low-depth sample and 75 copies for the high-depth sample. To evaluate whether depth was responsible for this discrepancy, we randomly subsampled the high-depth sample to ∼12.5X and estimated the copy number 100 times. The results ranged from 74 to 77 copies, indicating that the variation between the two replicates was likely not due to the sequencing depth, suggesting that experimental steps preceding the informatics are likely the source of the modest discrepancy exhibited by our estimates.

In the GDL, half of the stocks showed copy numbers between 90 (Q1) and 112 (Q3), roughly consistent with reports from the literature (100-111) (Lifton 1978; Strausbaugh and Weinberg 1982). Notably, half of the estimates were outside those boundaries, suggesting substantial natural variation in histone copy number. The lowest copy number estimate we observed (29) exceeds the experimentally derived floor for copy number (20) required for normal development (Zhang et al. 2019), and the highest (146) exceeds the typical copy numbers of 100-110 by more than 40%. Though the numbers we report are only estimates of the true number, the error is unlikely to explain most of the variation in our observations. The discrepancy between estimates based on assembly and Illumina read depth for iso-1, and A4 was about 10%, while the range of the estimates of the two replicates for ZW155-ST/ST was also about 10% (74-81). Of course, any individual experiment in a sample could exhibit dramatic anomalies due to unknown experimental errors. However, the rates of such errors would have to be quite high and affect only the coverage outliers to explain most of the variation we observe.

#### Comparative genomics of the histone cluster

Figure 3A shows a schematic view of the histone locus across 4 different assemblies: iso-1 (Flybase), iso-1 (HiFi), A4, and A3. The iso-1 FlyBase assembly contains a gap represented by empty space and is scaled to match our assembly’s 113 copies. The black box in the A3 assembly separates the left part (∼36 units) and the right part (∼61 units) of the assembled cluster. The light blue boxes represent different LTR TE insertions in the locus. Two main length variants of Histone cluster units in the Drosophila melanogaster are 4.8 kb and 5 kb (Colby and Williams 1993). The 5kb repeat unit has a 242-bp insert between H1 and H3 genes. These variants are present in a number of *Drosophila* species, indicating that their origin predates the speciation event (Strausbaugh and Weinberg 1982). Figure 3B shows the distribution of 5 and 4.8 kb types in the assemblies. In the 3 strains, the 5kb repeat significantly outnumbers the 4.8kb one, and they are generally clustered separately, with 4.8kb found exclusively towards the proximal flank of the array.

**Fig. 3:**
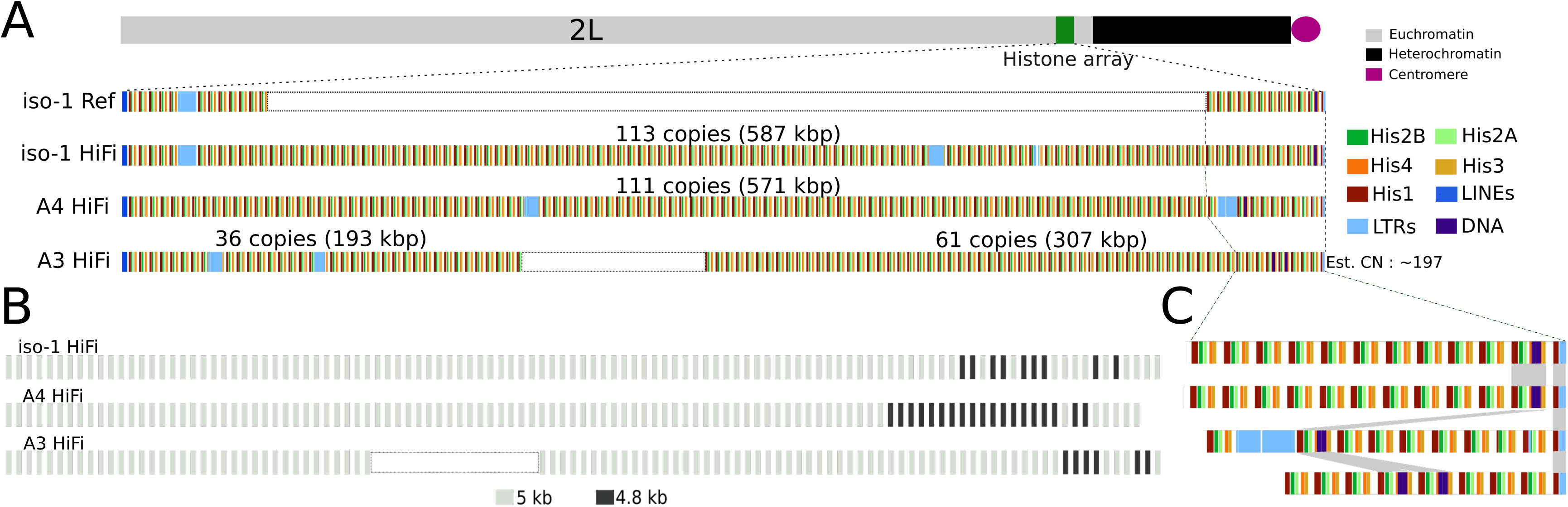
Maps of the newly assembled histone locus in Drosophila melanogaster. **A.** Overview of the structure of the histone locus. Top: Schematic illustration of the location of the histone locus relative to the rest of chromosome 2L. Bottom: Map of location of elements in the histone cluster, including the 5 individual histone genes and various transposable elements. “iso-1 Ref” refers to the Release 6 community assembly and “iso-1 HiFi” indicates the assembly presented here. White rectangles for iso-1 Ref and A3 HiFi indicate gaps resized to facilitate comparison with the other two assemblies. **B.** Location distribution of the major histone unit types (i.e., 5kb and 4.8kb). **C.** Expanded map of the proximal end of the array. Order of strains follows panel A.

Figure 3C shows an expanded version of the proximal end of the array. Given how the relative location of the *DNA/POGO*-containing unit varies within the array, the array has clearly expanded and/or contracted since the haplotypes in our sample last shared a common ancestor*. To* better understand the array dynamics, we sought to align the arrays and possibly comment on the mutational mechanism active in the array.

#### Anchored comparisons between highly repetitive arrays

To assess variation in the structure of arrays between different strains, we compared segments flanked by putatively unique anchor sequences. Even though the repeat units are too similar to each other to construct reliable alignments that are biologically meaningful, comparing haplotype segments flanked by the same anchors in different strains permits us to establish a conservative floor for estimates of structural variation. By comparing various metrics (e.g., the number, types, and arrangements of units) in these segments, we can determine whether single duplication or deletion events explain these changes or if the changes are more complex. Units with unique features that occur only once in each haplotype in a given comparison can serve as unique landmarks in the array we call anchors. This is the same approach used above for *DNA/POGO* TE-containing units. For example, when a TE is inserted into the array, the combination of the specific insertion location and TE type establishes a landmark that can be used as an anchor, provided it has not spread widely throughout the array through duplication and/or gene conversion. From this type of anchor landmark alone, we can already determine that the histone cluster experiences rapid expansions and/or contractions. The red boxes in Fig. 4A indicate such landmarks and occupy substantially different positions in the cluster between strains. Here, the TE insertion serves to mark the unit. However, any other reliably identifiable mutation unique in the array (e.g., deletions, insertions, or even unique SNPs) can serve as candidates for landmark anchors.

**Fig. 4:**
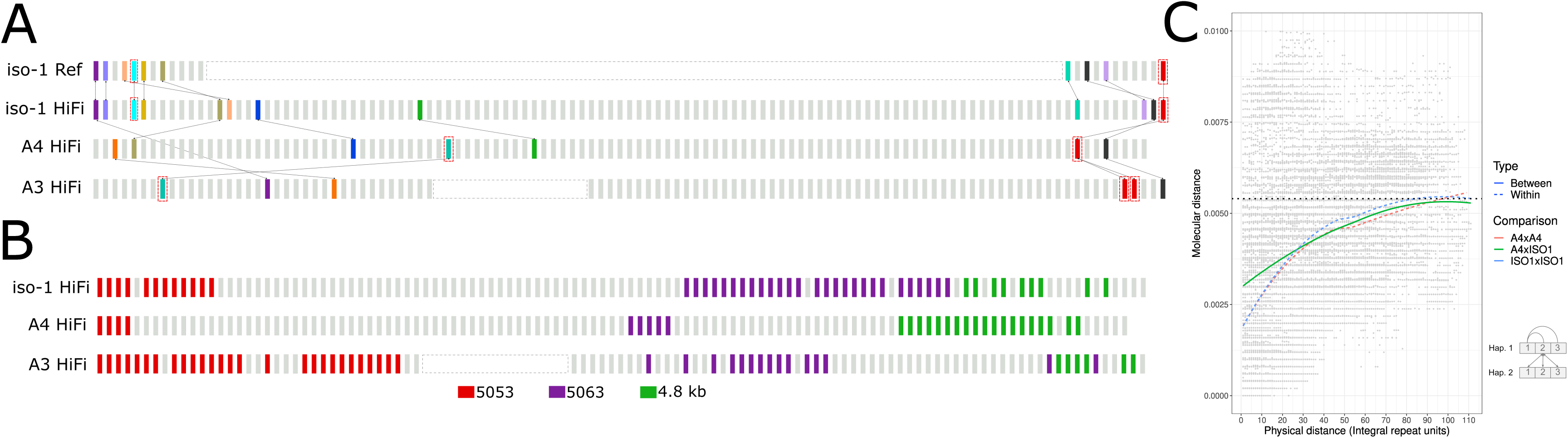
**A.** Distribution of landmark anchors used in this study. Anchors enclosed in red boxes indicate units harboring TE insertions. Other anchors are identified by phylogenetic grouping (see Methods). Anchors sharing the same color are putatively homologous. Lines between strains highlight the positions of these putative homologs in their respective haplotypes. **B.** Distribution of unit types along the histone cluster in 3 haplotypes, including two additional subtypes of 5kb: 5053 and 5063. **C.** Graphical representation of physical distance between histone units on the chromosome (x-axis) and molecular distance (y-axis). Molecular distances represent all unique pairwise comparisons either between or within strains. Bottom right: Schematic diagram how molecular distance comparisons between and within strain comparisons were conducted.

To systematically identify more putative anchors, we extracted individual units in the arrays for iso-1-Rel6, iso-1, A4, and A3. For every pairwise comparison, we drew a neighbor-joining tree for all units in both strains (500 bootstrap replicates, 50% cutoff for consensus tree). When two units from two different strains are clustered together, we mark them as unique anchor points. TE-containing units were analyzed separately, as described above. In addition to the unit containing *DNA/POGO*, one unit harboring HMSBEAGLE_I insertion was present in A4 and A3 but not iso-1. Figure 4A summarizes the structure of the histone units in relation to the anchors. Such putatively orthologous copies are marked using the same color. The TE-containing anchors are marked by red boxes. The sequence flanked by the same adjacent anchors in different strains is often dramatically different in size and/or composition, indicating a rapid turnover of repeat units in the cluster with units being added and deleted. This is true even for alternate assemblies of the same strain (n.b., these alternate assemblies are based on data collected in different years in different laboratories). In the relatively small fraction of the histone cluster present in the iso-1 Rel6 reference, the structure is not identical to the corresponding segments in our iso-1 HiFi assembly. The periphery of the histone locus was first added to the assembly in Release 4 (https://www.fruitfly.org/annot/release4.html). If the assembly of these regions in FlyBase is correct, it suggests that rapid evolution has occurred since iso-1 DNA was collected for the FlyBase reference and when it was collected for this study. This observation mirrors similar observations regarding structural variation in a previous study (Solares et al. 2018). In that study, several TE insertions and copy number changes affecting a 207-bp tandem array in *Muc26B* distinguished the two assemblies, indicating fast evolution.

#### Variation between histone units within and between strains

Histone repeat units are very similar to one another. To quantify the structure of this homogeneity, we annotated the distribution of various unit types across the cluster. To identify closely related repeat units, we focused on clades in pairwise phylogenetic trees composed of units from both strains. As expected, the 4.8 kb units are distinct from the other units, clustering together. We also identified two more unit types, 5053 and 5063, based on the median length of the identified units in the clades. Figure 4B shows the spatial distribution of these three broad unit types in the three strains. Two observations stand out: 1) the different unit types are present in different copy numbers in different strains; 2) the unit types tend to cluster together.

Next, we compare the similarity between histone units and their positions along the array, both within and between strains. Figure 4C shows the relationship between physical distance (x-axis) and molecular (i.e., evolutionary) distance (y-axis) between units in the array. For adjacent units in the same strain, the average molecular distance averaged within both strains is 0.12% (range: [0.112, 0.131]), more than tripling to 0.43% (range: [0.41, 0.46]) after about 30 units. However, when two units occupy adjacent positions in different strains, their average divergence is about 2.5-fold higher, or ∼0.31%, increasing to 0.41% after about 30 units. For longer separations, comparisons both within and between strains slowly asymptote from ∼0.41% when 30 units apart to ∼0.55% when 90 units apart. We can also examine individual differences between units instead of the average for a given physical distance. In this context, the maximum single difference we observed between any two units within the species is about 1.2%, which is between a 4.8kb and a 5kb unit. Within 5kb units, all comparisons differ by less than 0.9%. When we compare a randomly chosen unit from iso-1 (Unit 18) in *Drosophila melanogaster* to a 4.6kb partial histone unit cloned from *D. simulans* (accession AB055959 (Tsunemoto and Matsuo 2001)), the aligned portion of the sequences differ by ∼6.7% without indels or by about 9.2% when indels are included.

#### Linkage disequilibrium (LD) in the histone locus

To measure LD across the histone locus, we used data from the Zambian lines from the *Drosophila* Genome Nexus (DGN) (Lack et al. 2015). We employed this sample population because it is derived from haploid embryos, thereby mitigating issues related to genotyping. Moreover, the Zambian strains show high levels of inter-strain variation. Ignoring the stocks with the *In(2L)t* inversion, 147 of 197 haplotypes were analyzed from the Zambian population.

Fig 5A shows IGV plots of two windows flanking the histone locus. We label the flanking SNPs in relation to the position of the histone cluster. Distal positions start with -1 and become more negative the more distal they are. Proximal positions start with 0 and become more positive the more proximal they are. Two major haplotypes span the region, which we call H1 and H2. For the sake of simplicity, we have neglected SNPs and indels with MAF < 5%. However, this does not impact our observations since they are in *complete LD* (D’=1) with their respective haplotypes. Out of 147 stocks, 145 are either H1 or H2. Only 2 samples are exceptions: ZI170 and ZI220.

**Fig. 5:**
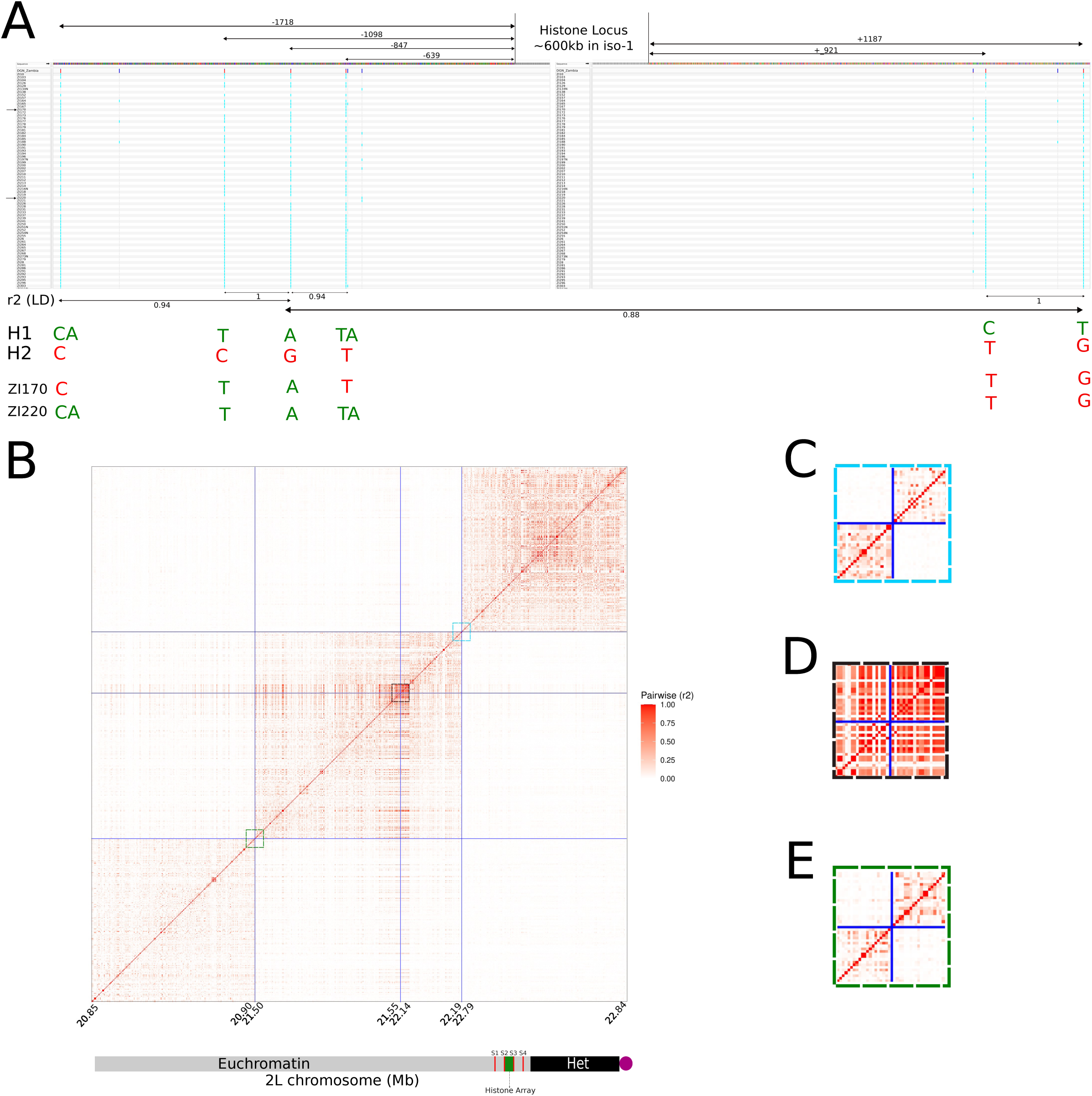
Recombination in and around the histone locus. **A.** Integrative Genomics Viewer plot of the regions flanking the histone locus. Values between double headed arrows under the x-axis indicate the r2 measure of linkage disequilibrium between the variants spanned by the arrows. Coordinates are with respect to the histone locus, with negative values indicating the distal flank and positive values indicating the proximal flank. H1 and H2 each represent haplotypes present in all but 2 strains. ZI170 and ZI220 possess their own private combinations of variants from H1 and H2 haplotypes and are the only representatives of their respective haplotypes. **B.** Matrix of r2 between 4 50kb segments (S1-S4) around the histone locus. Each segment is fixed in physical size. Since only variants are plotted, the plotted size scales with the number of variants in the segment. Each segment is separated by a size determined by the length of the histone cluster in iso-1 (600kb). Colored boxes correspond to expansions in panels C-E. Bottom: Schematic representation of chromosome 2L and the location of the histone locus with respect to the segments and to the annotated boundaries of the heterochromatin. **C.** Expanded view of pairwise LD for 50kb regions flanking a control locus spanning 22.19Mb-22.79Mb (i.e., S3 and S4). **D.** Expanded view of pairwise LD for 50kb regions flanking the histone locus spanning 21.55Mb-22.14Mb (i.e., S2 and S2). **E.** Expanded view of pairwise LD for 50kb regions flanking a control locus spanning 20.9Mb-21.5Mb (i.e., S1 and S2).

The first variant on the distal flank is at position -639 bp. The proximal end of the histone cluster terminates in a partial unit of about ∼1.5 kb in all 3 strains for which we have the assembly. The first ∼1kb of the proximal flanking region is repetitive and exhibits sparse read coverage. Some SNPs called in this region exhibited elevated rates of missing read data for some strains and had lower-quality mapping scores. For example, SNP +921 had 3.4% missing data and an average MAPQ of 43 (compared to almost 60 elsewhere). The r^2^ between the first high-quality SNPs in each flanking region (positions -847 and +1187) is 0.88. In this comparison, while we neglect SNP +921, it is in *perfect LD* (r^2^ = 1) with SNP +1187. Except for only two of the strains mentioned above (i.e., ZI170 and ZI220), the entire collection is in *perfect LD* (r^2^=1) between -847 and +1187.

Fig. 5B shows pairwise LD on a larger scale. The blue lines demarcate four 50kb segments. Because the plot depicts pairwise r^2^ between sites, it is only defined between two variable sites. As a result, the different sizes of identically sized segments in the plots reflect different SNP densities in those segments. Each segment is separated by ∼600kb (the size of the histone cluster in iso-1) in positions indicated by the blue lines. The actual histone cluster separates segment pair 2:3, whereas pairs 1:2 and 3:4 are separated by other sequences downstream and upstream of the histone locus, respectively. LD typically decays rapidly in *D. melanogaster*, so two variants separated by ∼600kb should not be in LD. That holds true for segment 1 versus 2 and segment 3 versus 4 (Figure 5C, E). However, segments 2 and 3 exhibit extremely high LD even across 600kb (Figure 5D).

### Structural variation in the *Stellate* clusters

#### Euchromatic *Stellate* cluster

Our assemblies permitted us to analyze the copy number of both X-linked *Stellate* clusters (Figure 6A). In the euchromatic *Stellate* cluster spanning bands 12E1-12E2, the three strains iso-1, A4, and A3 exhibited three very different configurations, with the most notable variation on the proximal end of the cluster (Figure 6B). Except for three LTR insertions in iso-1, the sequence flanking the distal side cluster is not structurally variable between strains, with all strains carrying a single copy of *Stellate (*called *Stellate 12D orphan*) preceding either ∼35kb (in A3 and A4) or ∼52kb (in iso-1) before the beginning of the *Stellate* cluster proper. The proximal end of the locus, on the other hand, shows significant variation. Our HiFi assemblies carry 11, 198, and 1 copy for iso-1, A4, and A3, respectively. While our assembly of iso-1 bears 11 tandem copies in this region, the rel6 assembly of iso-1 shows 12 copies. The plot of HiFi read depth supports the locus structure of *Stellate* in our HiFi assembly, showing no hallmarks of collapse or misassembly in the read depth plot (Supp Fig 16A). Additionally, the arrangements of repeat units in the cluster differ between iso-1 HiFi and iso-1 rel6, similar to our observation in the histone cluster (Supp Fig 17). While this may indicate an error in the public assembly based on older data, it’s also possible that this reflects the rapid evolution of copy number between the parental strain and its derivatives sequenced years later (Solares et al. 2018). Finally, in addition to carrying 198 tandem copies spanning ∼260kb, the proximal portion in A4 carries a ∼9kb ROO insertion. Like iso-1, the read depth plots for A4 and A3 show relatively uniform coverage (Supp Fig 16B, C).

**Fig. 6:**
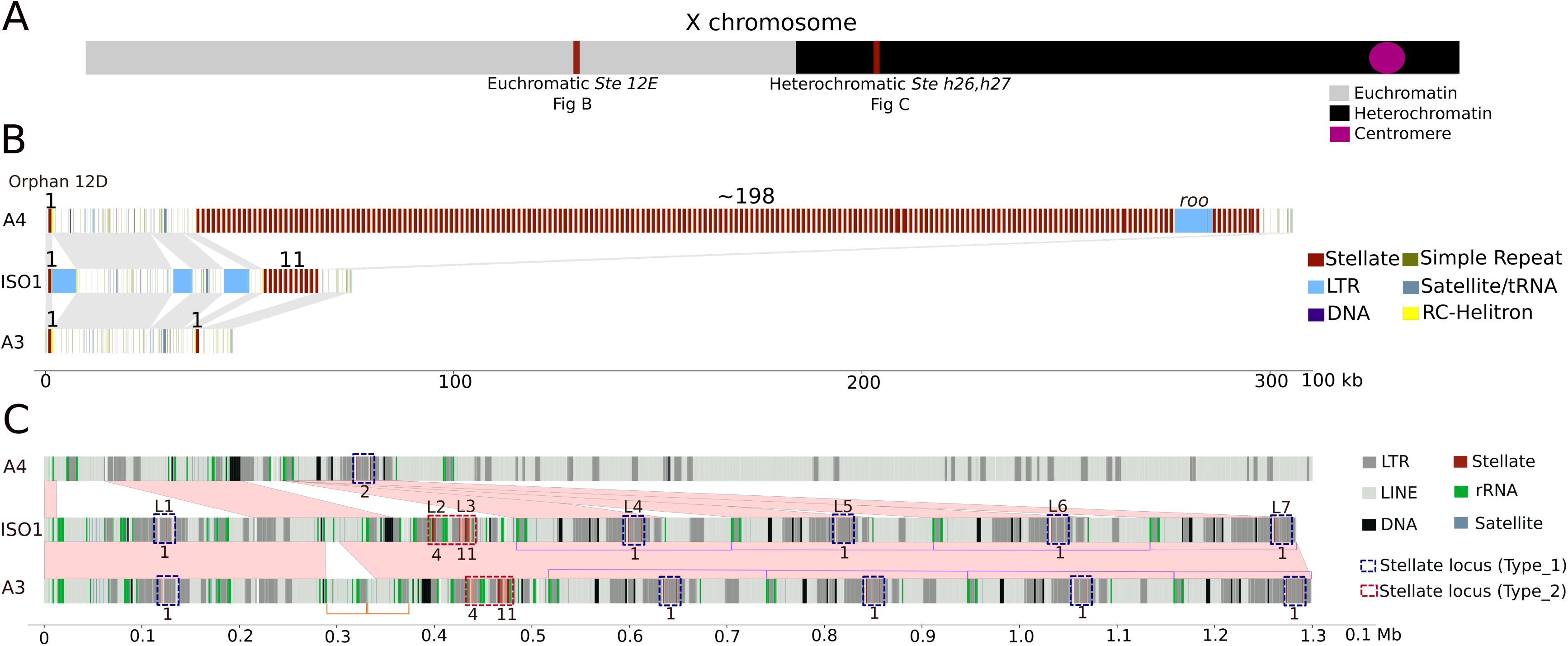
Maps of euchromatic and heterochromatic *Stellate* clusters. **A.** Schematic illustration of the location of the Stellate clusters relative to the rest of the X chromosome. **B.** Overview of the structure of the euchromatic Stellate locus. The numbers on the top represent full length *Stellate* sequences in the tandem array. **C.** Overview of the structure of the heterochromatic Stellate locus. Type 1 and Type 2 loci are marked by labels on the top in iso-1. The numbers on the bottom represent full length *Stellate* sequences in the particular locus. The skin color parallelograms represent syntenic sequences (drawn by eye). Purple half rectangles represent the 200 kb tandem segmental duplicates harboring some of Type 1 loci.

#### Heterochromatic *Stellate* cluster

The heterochromatic *Stellate* cluster (*hetSte*) spans the *h26* and *h27* regions of the X chromosome (Tulin et al. 1997). This cluster is part of the newly assembled X-linked sequence (see Fig 2B, C). We distinguish between two major cluster subtypes containing copies of *Stellate*. The first (labeled Type_1) contains a single full-length copy of *Stellate*. It appears to be derived from a tandem array of four 1150-type repeat copies that acquired various insertions and deletions, leaving only one stellate copy intact. The second (Type_2) contributes the most copies to the X and comes in two forms, one with 4 and one with 11 intact tandem copies (Fig. 6C).

The iso-1 strain carries 5 loci of the Type_1 subtype, including the most distal locus (L1) and the 4 most proximal loci (L4-L7; Fig 6C). L1 contains 4 copies of *Stellate* (3 partial and 1 complete). In addition to *Stellate*, this sequence contains insertions of *LINE/R1*, *MDG1-LTR* and *NINJA-LTR* (Supp Fig 18). In this L1 variant of the Type_1 subtype, the full-length *Stellate* copy is present in the first unit. The L4, L5, L6, and L7 loci are also Type_1 derivatives, but the first unit is missing, and the second element lacks the small *LINE/R1* sequence (Supp Fig 18). In contrast to L1, for L4-L7, the full copy of *Stellate* is present in the 3rd unit. All Type_1 sequences possess a *NINJA* insertion at the 90th position of the 3rd unit, though the L1 locus also carries a partial deletion of *Stellate* (Supp Fig 18). Thus, all Type_1 loci in iso-1 possess only one full-length *Stellate* repeat. The L4-L7 loci are within a series of tandem segmental duplicates, each ∼200kb in length, and are present on the opposite strand relative to L1 (purple rectangles Fig 6C).

The L2 and L3 loci are Type_2 and comprise the main tandem *hetSte* clusters. They consist of 5 (4 complete, 1 partial) and 12 (11 complete, 1 partial) *Stellate* repeat units respectively (Fig 6C). Each locus comprises a tandem *Stellate* cluster flanked on one side by MICROPIA and on the other by BATUMI TEs and a mix of R1/R2 transposons and rDNA sequences, which we abbreviate as mSbr. The L2 and L3 loci are adjacent to each other in an inverted tandem orientation (i.e., [mSbr][rb’S’m’], where the ‘ indicates the opposite orientation) (Supp Fig 19). As a consequence, the *Stellate* clusters in these loci are separated by ∼24 kb of sequences made up of BATUMI, rDNA and R1/R2 and are on opposite strands (Supp Fig 19). Given that the first unit of both clusters contain the same partial deletion and that the clusters are on opposite strands, they are likely the result of tandem inversion duplicates (TIDs) (Reams and Roth 2015). Similarly, Type_1 loci L1 and L4, L5, L6, and L7 are also flanked by BATUMI and R1 sequences and are on opposite strands, again suggesting that these are TIDs. A3 bears the same haplotype as iso-1 (except for a small duplication (orange rectangles Fig 6C)) in this region and hence has the same copy number. A4 exhibits a dramatically different structure in this region (as previously described in Fig 2C) and has only one Type_1 locus similar to L4, L5, L6, L7 (Fig 6C). However, in this case, the first element is not deleted, so it has 2 full-length copies of the 1150-type repeat.

#### Comparative Genomics of *Stellate*

To compare tandem units of *Stellate* clusters, we followed a similar approach to our comparisons of the Histone cluster. Briefly, our approach relies on using anchors of homology-based on diagnostic differences that make a repeat unit unique within a strain. In the euchromatic cluster, we identified only three landmark anchors in iso-1 and A4, two of which are the most distal and proximal units in both strains (Supp Fig 20). The most proximal unit is missing the final 302 bp of the repeat. In A3, this is the only one present. In the heterochromatic cluster, A3 and iso-1 exhibit the same number of repeats in the L2 and L3 loci. In fact, the L2 locus (5 copies) exhibits the same structure in both strains (Supp Fig 21A). The structure of the L3 locus (12 copies) is relatively stable. When identifying landmark anchors, we find that most units are reciprocally most related to another unit in the same position in the array, with one exception: a 1 unit insertion/deletion in both homologous clusters and a single mismatched unit (Supp Fig 21B).

A recently published heterochromatin-enriched drosophila assembly (Chang and Larracuente 2019) (now referred to as *iso1_HetEnr*) also reconstructed heterochromatic stellate sequences. We discovered the L1 locus in *Contig_135* (440 kb). What we call L2 and L3 bearing the 5 and 12 copy tandem arrays of *Stellate* are found in their *Contig_5* (535 kb). The structure of L2 and L3 in *iso1_HetEnr* matches exactly (to the base pair) with our HiFi assembly, with variants in both L2 and L3 having a one-to-one correspondence with *hetSte* variants in our HiFi assembly. In a small 35kb contig called *Contig X_9*, we also located sequences corresponding to the L4-L7 *Stellate* loci. The sequence in *ContigX_9* spans the *Stellate* sequences and some flanking sequences on both sides of *Stellate* that corresponds to a subset of the ∼200kb unit of the 4-unit tandem array in our assembly (Fig 6C). We also inspected the tandem euchromatic cluster in *iso1_HetEnr* and found a total of 12 copies. This assembly represents a third assembly of the euchromatic cluster from the iso-1 strain. Perhaps unsurprisingly, it exhibits a different structure than observed in the Rel6 or our HiFi assemblies (Supp Fig 17).

## Discussion

### Routine assembly leads to improved reference genomes

The assemblies of three near-isogenic *D. melanogaster* strains generated here exhibit exceptional levels of completeness, contiguity, accuracy, and low levels of heterozygosity (i.e., single copy BUSCOs 99.27-99.42%, contig N50s of 21.5-23.0 Mb, base level Phred 50-56, and heterozygosity 0.07-0.08%). Notably, our assembly of a descendant of the reference strain iso-1 added an additional ∼8.0 Mb of sequence to the autosomes and X chromosome arms compared to the latest release (R6) of the reference genome assembly. This is almost half (∼47%) of what was added to those arms between the initial sequencing of the *D. melanogaster* genome (R1) (Adams et al. 2000) and the most recent version (R6) released in 2014 (Hoskins et al. 2015). Release 1 recovered ∼66% (116.2/175 Mb) of the estimated female genome (Bennett et al. 2003; Ellis et al. 2014) in chromosome arm scaffolds in 2000 (Adams et al. 2000). By 2014, Release 6 improved this by about 10% (to ∼76%, or 133.0 Mb), including scaffolds and auxiliary sequences (Hoskins et al. 2015). Considering only primary chromosome scaffolds, Release 6 placed only ∼127.5Mb, or ∼73% of the genome. In comparison, we independently recovered ∼81% of the genome (141.9 Mb) in primary chromosome scaffolds alone, representing an improvement of 8.2%. Such improvements are consequences of the long (median 13.7-15.6 kb) and accurate (average ∼Q28) reads used in the assembly. Importantly, these assemblies reliably render previously recalcitrant regions tractable for study.

The most substantial improvements to our assemblies have been in the most repetitive regions of the genome, particularly the heterochromatin. Notably, these improvements relative to previous long-read approaches (Chakraborty et al. 2018; Chakraborty et al. 2019) mostly result from improved accuracy. For example, in the reference strain, we extended the proximal end (i.e., the end nearest the centromere) of the X chromosome by ∼3.21 Mb relative to Release 6. This heterochromatic region also contains the initial ∼450 kb of the rDNA array, which is estimated to be ∼2.8 Mb (200–250 rDNA units) (Bianciardi et al. 2012). Our assemblies of A4 and A3 performed similarly, recovering these same regions and ∼522 kb and ∼1Mb of the rDNA, respectively. In contrast, Release 6 (Hoskins et al. 2015) reports only ∼75kb of auxiliary scaffolds for rDNA. We also identified 15.7 Mb of putative Y-linked sequence by using only routine assembly procedures in our reference HiFi assembly. This more than quadruples that reported for Release 6 and exceeds even that of (Chang and Larracuente 2019) by ∼1Mb, who employed a clever and labor-intensive approach based on enriching sequencing reads for heterochromatin and assembling them with parameters most suited for such repetitive regions followed by manual curation.

### Newly assembled X-linked heterochromatic sequence

The rDNA locus and the surrounding sequences in X pericentric heterochromatin are highly repetitive and have resisted reconstruction at the sequence level. In arthropods, in addition to rRNA genes, the rDNA region also harbors two TEs unique to the rDNA locus called LINE *R1* & *R2*. *R1* and *R2* possess specific insertion sites only in 28s rDNA of arthropods (Pérez-González and Eickbush 2002). *R1* is especially abundant in the distal region flanking the rDNA, occupying almost ∼50-60% (Fig 2C). Given its target site specificity, so many *R1* TEs in the region are unlikely to represent solely *de novo* insertion events. Instead, TEs can also form tandem repetitive arrays driven by unequal exchanges (McGurk and Barbash 2018). However, the newly assembled sequence does not resemble a standard homogenous tandem array, exhibiting a high degree of heterogeneity. In repetitive regions such as these, higher-order repeats (HORs) sometimes arise, driven by unequal exchanges (Suzuki et al. 2020). HORs may exhibit different organization either because they originated independently or because they acquired various deletions and insertions independently over time (such as other TEs). Our observations posit a model where a region primarily containing *R1* tandem elements arose in the rDNA locus. Subsequent cycles of amplifications (and deletions) and the emergence of new HORs, driven by unequal exchanges, could result in the heterogeneous structure we observe today. The dot plots of the region (Supp Fig 6) confirm the nature of tandem repetitive content reminiscent of dot plot patterns encountered for centromeric repeats containing HORs. And the variable copy number segmental duplicates (Fig 2A) likely represent larger-scale HOR structures. Using short-read data, (McGurk and Barbash 2018) estimated copy number variation within the X-linked *R1* tandem array and used unique junctions to infer the heterogeneity of the array. However, with our fully resolved map of genetic variation within the locus, not only can we infer that these arrays are quite different, we can propose models to explain the mutational origins of the entire region. However, due to the absence of any apparent functional sequences, we cannot infer whether or not adaptation plays a role in the evolution of this region.

Due to the highly mutable nature of the structure of such loci (Nurminsky et al. 1994; Campbell and Eichler 2013) and the large effective population size of *D. melanogaster*, individual haplotypes are likely to exhibit large structural differences from one another, including sometimes dramatic variation in copy number. Thus, it is possible that the mutation underlying the ∼250kb segment collapsed in our iso-1 assembly (Supp Fig 7) is a very recent expansion. Such recent expansions would create two virtually identical segments, making them hard to resolve even with long, accurate reads (Porubsky et al. 2023).

### Structural variation in the histone cluster

#### Assembly of *D. melanogaster* histone cluster for iso-1, A4, and A3

Until recently, the precise structure of the histone cluster was virtually impossible to reconstruct. The histone locus consists of a tandem array of ∼100-110 tandem units of a ∼5kb sequence Each repeat unit contains the 5 histone genes nearly identical to the other units. Though the histone cluster is large and homogeneous, some variation exists between the units (Fig 3-5). Despite this variation, the long stretches of virtually identical repeats within the cluster have rendered the histone locus a substantial challenge to assemble. The most reliable DNA sequencing-based model of the structure histone cluster was painstakingly reconstructed via a bespoke assembly approach that takes advantage of machine learning (P. Bongartz and Schloissnig 2019). Such approaches target identifying the comparatively weak signal (actual sequence variants that enable read overlapping) in datasets replete with noise (the high sequencing error rate of long read sequencing approaches). With recent improvements to error rates in long-read sequencing (e.g., from 1 error in 10 bp to 1 error in 1,000 bp per sequencing read (Marx 2023)), we can now accurately reconstruct the histone cluster with routine application of standard assembly approaches (Supp Fig S8-9, S11-12). Our results were virtually identical to those of (P. Bongartz and Schloissnig 2019). Such remarkable similarity between independent sequencing datasets and alternate assembly approaches gives us confidence in our assembly and any analyses we do further downstream. The apparent disagreement in the most proximal 2 units suggests either assembly error in the i*so1-BS* genome or a polymorphism segregating in the strain, as we identified individual reads supporting our assembly model. Even though we did not have to resort to novel assembly software to solve the locus, our approach did require targeted assembly of the histone locus with parameters appropriate for extremely repetitive sequences (see Methods) and was not successful for every dataset (Supp Fig S13-14). Nevertheless, these improvements in the reconstruction of the histone cluster enable an unprecedented view into the structural variability in this segment of the genome encoding the proteins necessary for nucleosome formation.

#### Dramatic variation in copy number and organization of histone repeat units

Our iso-1 and A4 assemblies indicate copy numbers of 113 (587 kb) and 111 (571 kb), respectively. However, the data from A3 did not yield a reliable assembly spanning the entire locus. In the initial assembly of A3, which indicated a cluster size of 114 copies, read mapping coverage to the histone cluster indicated a misassembly and collapsed regions (Supp Fig 13) . The standard targeted assembly did not resolve the locus, but it suggested the size of the histone cluster to be at least 134 total units (spread across 3 contigs). An alternate approach to estimate the copy number based on the depth of Illumina read coverage estimates the size of the A3 histone cluster to be 197 units. Thus, based on estimates ranging from 134 to 197 units, the size of the histone cluster in A3 is between 21% and 77% larger than in iso-1. Both estimates are substantially higher than expected from previous surveys of the histone locus, which typically converges on a copy number of roughly 110 copies (Lifton 1978; Strausbaugh and Weinberg 1982).

These results suggest dramatic copy number variation in the histone cluster may be common. To independently validate this, we used Illumina read coverage in the *Drosophila* Global Diversity Lines (GDL) (Grenier et al. 2015). Our survey of 85 GDL strains resulted in a median of 104 copies (Q1=90 and Q3=112), which is in line with other estimates. Surprisingly, across the entire dataset, the copy number varied over 5-fold, ranging from 29 to 146. Conservatively, this is strong evidence for several-fold natural copy number variation in the histone locus. Despite this wide variation (especially the low copy number of 29), our observations are consistent with experimental evidence in histone knockout lines, which suggests that transgene copy numbers as low as 8 are viable and as low as 20 restores near wildtype fitness (Zhang et al. 2019). Additionally, the RNA or protein levels between flies with either 12 or 20 copies show no significant difference (Zhang et al. 2019). This indicates dosage compensation mechanisms are active in histone protein production, possibly leading to relaxed selection on copy number.

Among our assemblies, the organization of the histone locus is also quite variable (Fig. 4A, B). To characterize this dramatic variation and elucidate the dynamics of the array, we used unique mutations (e.g., TEs) within the cluster as reference points (i.e., landmark anchors) for determining homology between repeat units. The major repeat unit types (5kb and 4.8kb) are similarly distributed across the locus, with the less common type (4.8kb) being concentrated at the proximal end of the locus (Figure 4B). However, even among the 3 strains we surveyed, the details differ substantially in terms of the copy number of the 4.8kb type and how it is interspersed with the 5kb type. We observed a wide variation in the copy number of the 4.8kb type between strains (9, 19, and 6 copies in iso-1, A4, and A3, respectively; Fig 4B). Additionally, the copy number and locations of the 5kb types within the 4.8kb-rich segments vary dramatically between strains (7, 1, and 3 5kb types interspersed inside the 4.8kb types in iso-1, A4, and A3, respectively; Fig 4B). A similar picture emerges when we study subsets of the 5kb type. There are two subtypes of the 5kb type we call 5053 (Figure 4B, distal end) and 5063 (Figure 4B, proximal end). Not only do the copy numbers of these subtypes vary considerably between strains, but the spacing between the units seems to be distinct between strains as well. Even more organizational diversity is evident when we consider the structure of the locus from the perspective of landmark anchors (Figure 4A). We can see that anchors vary not only with respect to their relative positions in the array but also that certain pairs of anchors can swap their relative orders (see the crossing of dotted lines in Figure 4A). Such variation is incompatible with simple expansion or contraction, requiring multiple mutational steps. Given that gene conversion and repeat expansion and contraction are ubiquitous in these regions, this opens up several possibilities. One scenario is the gene conversion of one anchor followed by the loss of the first copy (either through another gene conversion event or through deletion). Another possibility involves a duplication followed by at least two deletions (Supp Fig 22).

#### Concerted evolution in the histone cluster

In our results, the histone cluster is extremely homogeneous, exhibiting at most a 1.2% difference between any two units of the array, both within and between stains. This variation is often dramatically lower but is typically between 0.12% for adjacent units and 0.55% between units on opposite sides of the locus for intra-strain comparisons. However, comparisons between *D. melanogaster* and *D. simulans* show a 6.7% divergence. This is 5.6-fold higher than the most divergent comparison (1.2%) within *D. melanogaster*, a classic signal of strong concerted evolution (Matsuo and Yamazaki 1989; Hickey et al. 1991). These results generalize previous conclusions concerning H3 variation (Matsuo and Yamazaki 1989), which showed an 8.5-fold difference. The repeat units in the histone locus exhibit stronger homogenization between units that are closer to one another, both within and between strains (Fig. 4C). Additionally, homogenization is higher within strains than between, though only within about 30 units (i.e., ∼150kb), after which the genetic variation between repeat units is determined primarily by their relative position in the array rather than whether they are found in the same strain (Fig. 4C).

#### Suppression of crossing over across the histone cluster

One surprising observation is the near absence of evidence of crossing over in the segments surrounding the histone cluster. In the 147 DGN strains, SNPs immediately flanking either side of the histone cluster are in complete LD for 145 of them (Fig. 5A). In the Zl170 strain, the pattern of variation in segment 2 (S2 in Fig. 5B) is consistent with either a double crossover event or a gene conversion event on the distal flank of the histone cluster. In Zl220, the polymorphism data is consistent with a crossover event occurring between the distal and proximal flanks of the histone locus (i.e., between S2 and S3; see Fig. 5B), though it is also consistent with gene conversion in the proximal flank (S3). In this case, ∼3 kb (0.5kb distal, 1.5kb partial unit, 1kb proximal) of flanking sequence plus the ∼600kb (depending on the strain) of the histone locus is between the verifiable changes in Zl220 haplotype (Fig. 5A). In both cases, the haplotype switch is embedded in a region of otherwise extremely elevated LD extending over hundreds of kilobases (Fig. 5D). This makes it more likely that the level of recombination we observe is a result of gene conversion rather than crossing over.

#### Concerted evolution is primarily driven by intrachromosomal recombination

Previous studies focused predominantly on interspecies comparisons due to the lack of full-length arrays in genome assemblies, limiting our understanding of the underlying mechanisms of concerted evolution. However, analysis of fully resolved arrays within a species establishes patterns occurring at shorter evolutionary timescales, which are relevant to understanding the mechanisms driving concerted evolution in tandem repeat clusters. The variation in copy number of broader unit types (Fig 4B) across different haplotypes and their propensity to cluster within single haplotypes implies that homogenization predominantly occurs through recombination within a single chromosomal lineage. This suggests that the rate of intrachromosomal recombination is significantly higher than recombination between homologous chromosomes (interchromosomal recombination) (Supp Fig 23). Gene conversion and crossing over during meiosis are two ways through which interchromosomal recombination may occur. While gene conversion in such tandem arrays is hard to assay, the rate of meiotic crossovers can be estimated using SNP markers flanking the arrays. Fig. 6 shows nearly no LD decay between markers separated by 600kb, indicating extremely low crossing over at or near the histone locus. Such findings align with previous observations of significant LD across tandem arrays in human rDNA (Seperack et al. 1988) and U2 snRNA (Liao et al. 1997). The observation that similar repeat types tend to cluster together (Fig 4B,C) also suggests that intrachromosomal events that are homogenizing repeat copies occur through patchwise local interactions (Durfy and Willard 1989).

Concerted evolution is thought to result from DNA recombination mechanisms, primarily gene conversion and crossover. Conversion is a secondary consequence of recombinational repair mechanisms. DNA damage, especially Double Stranded Breaks (DSB), is dangerous and can lead to genomic instability if not repaired correctly. Homologous Recombination (HR) is one of two major DSB repair pathways. HR may result in unidirectional transfer of information from one DNA molecule to another (i.e., gene conversion) and/or reciprocal exchange between two DNA molecules (i.e., crossover). In HR repair, the homologous template can be either the homologous chromosome, sister chromatid, or, in the case of highly repetitive sequences, loci on the same molecule. **(A)** Sister chromatid exchange and **(B)** intramolecular exchange are two ways through which intrachromosomal HR is realized. **(A)** During the G2/S phase in mitotically dividing cells, sister chromatids are preferred for repair, involving gene conversion and/or crossover (Kadyk and Hartwell 1992; Johnson and Jasin 2000). Unequal Sister Chromatid Exchange (USCE) can result in copy number changes via duplications and deletions in tandem repeats, similar to what we observe in the Histone cluster (Tartof 1974; Axelrod et al. 1994). Additionally, Long Tract Gene Conversion (LTGC) events between sister chromatids can produce gene amplifications (Johnson and Jasin 2000; Puget et al. 2005) relevant to the histone locus. **(B)** Intrachromatid recombination (i.e., intrachromatid gene conversion and intrachromatid crossing over) within tandem repeat arrays may lead to homogenization and copy number variation (Cohen et al. 2003), suggesting a role for intramolecular mechanisms in the rapid turnover and diversity of unit structures observed.

Our analysis, consistent with previous studies (Seperack et al. 1988; Schlötterer and Tautz 1994; Liao et al. 1997), indicates that intrachromosomal exchanges, potentially including USCE, LTGC, and intramolecular gene conversion and crossing-over, are the primary candidates for mechanisms driving concerted evolution at the Histone locus. While the focus is often on meiotic events as a source of heritable variation in organisms, mitotic events occurring in precursors to germ cells also potentially contribute to patterns we see in concerted evolution. Despite the dominance of intrachromosomal recombination in the evolution of tandem repetitive loci, sufficient interchromosomal recombination is necessary for maintaining similarity among repeat copies within a species relative to sister taxa. Occasional gene conversion (without reciprocal exchange) between homologous chromosomes must occur to mediate interchromosomal recombination (Liao et al. 1997).

### Structural variation of the X-linked *Stellate* clusters

Our results show that most copy number variation for *Stellate* stems from the X-linked euchromatic cluster (*euSte*), confirming (Palumbo et al. 1994) conclusion. Indeed, if we exclude the orphan copy in 12D, we observe variation from as few as 1 copy in A3 up to ∼198 in A4, with iso-1 harboring 11 copies (Figure 6B). Our estimate from iso-1 is fairly consistent with estimates from other assemblies of the strain, including iso-1 *HetEnr* and iso-1 Rel6, exhibiting 12 copies. However, all versions exhibit differences in their organization (Supp Fig 17). Such differences can be attributed to mutational mechanisms active in tandem arrays, though misassembly of either assembly in comparison could also be the cause. These mutation types are special in that they may accumulate relatively rapidly and result in dramatic changes in locus organization, especially compared to our expectations from SNP and low indel mutation rates of ∼1×10^-9^. Another cluster of *Stellate* genes resides in the heterochromatin (*hetSte*) of the X chromosome. We classified two major cluster subtypes of *hetSte* based on the organization of *Stellate* repeat units: Type_1 and Type_2. Type_1 is derived from a 4-unit tandem array, which acquired various indels, leaving only 1 or 2 stellate copies intact. The homogenous tandem arrays of *hetSte* repeat units were designated as Type_2. Since Type_1 loci have only 1 or 2 full-length copies in the three strains we studied, they might not be the main contributors to copy number variation in *hetSte*. Type_2 loci with their tandem arrangements might be responsible for the lion’s share of variation in *hetSte*. Type_2 loci (L2, L3) exhibit very few differences in arrangement between iso-1 HiFi and A3 and no difference between our iso-1 HiFi and iso-1 *HetEnr* assemblies. This contrasts the euchromatic cluster, where even the same strain iso-1 exhibits rapid turnover of these units for a similar number of repeat units. Failure to find many differences between A3 and iso-1 indicates that the tandem *hetSte* evolves slowly, which might explain the limited variability in this cluster.

Previous studies of *hetSte* have inferred independent origins of the euchromatic and the heterochromatic *Ste* clusters (Chang and Larracuente 2019; Adashev et al. 2020). We used fully resolved sequences of *hetSte* to show that all Type_1 and Type_2 *hetSte* loci can trace their origin to a single ancestral array, duplicated through either tandem inversion duplicates (TID) or being part of larger segmental duplicates. These segmental duplicates are examples of the HOR structures we described for the rDNA distal X-linked locus. Upon further inspection, we find that even Type_1 and Type_2 have common flanking sequences derived from BATUMI TEs on one end, pointing to a possible single origin of the heterochromatic clusters. Similarly, the first stellate orphan sits on the opposite strand with respect to the main *euSte* tandem cluster. Both loci have fragments of DINE_1 TEs plus a very small partial stellate fragment flanking one of the sides. This again suggests that the two loci might have originated from a TID event, and subsequently, one of them amplified. This is similar to the conclusions reached by (Kogan et al. 2012) (though they implicate DINE-1 as a driver of duplication). Hence, our analyses based on the detailed maps of these regions suggest that both the euchromatic and the heterochromatic clusters have originated via single independent copying events and subsequently amplified under genetic conflict.

#### Concerted evolution in Euchromatic stellate cluster

The *euSte* cluster exhibited extreme variation in copy number (from 1-fold to 198-fold), prompting further investigation into concerted evolution at the *Ste* clusters. The relationship between physical distance and molecular distance between repeat units for A4 *euSte* shows a different pattern compared to the histone cluster. Adjacent copies are not more similar than separated copies (Supp Fig 24A). Several obvious explanations fall short of explaining this pattern. Rapid array-wide homogenization could lead to such a uniform pattern. However, we do not see any significant differences for within-versus between-haplotype comparisons, as expected in the case of rapid homogenization (Supp Fig 24B). While the recent emergence of the *Stellate* array compared to the histone array would have afforded less time to accumulate variation between *Stellate* units than Histone units, we nevertheless find similar levels of variation (median of all pairwise distances) in Stellate (0.0031) and histone (∼0.004) clusters. In contrast to the histone cluster, if the relative rates of intra- vs inter-chromosomal exchanges are similar in the *Stellate* cluster, we also might see no distance-dependent effect. Additionally, the size of a single unit, the total size of the entire cluster, the possible linear range of localized interactions for gene conversion/duplication events, and the median length of such events – all these interdependent factors taken together can also contribute to the observed difference. The availability of more full-length *euSte* clusters would allow us to dig deeper, and it is left for future work.

### Persistent limitations in mapping the dark matter of the repetitive genome

While the work reported here is substantial progress compared to previous work, the heterochromatic regions we discussed illustrate some of the challenges that persist. Indeed, in the human telomere-to-telomere project, the rDNA is one of the only remaining frontiers where sequencing does not solve the region (Nurk et al. 2022). In our case, in addition to not assembling through the entire rDNA array, there are clear problems both adjacent to and inside the rDNA region. Specifically, the elevated sequence read coverage in that region strongly suggests that it has been collapsed in the assembly (Fig. S6). We also recovered an additional 6.4 Mb of rDNA sequences in our iso-1 assembly; 4.26 Mb was assigned to X, 1.48 Mb to Y, and 0.67 Mb was left unassigned. We could not order and orient these sequences, so they remain spread across 29 contigs, complicating downstream variation analyses. The assembly and scaffolding of the entire Y chromosome face similar challenges. While the iso-1 assembly did largely recapitulate results from a specialized approach employing targeted enrichment of heterochromatin for reconstructing the Y chromosome (Chang and Larracuente 2019) (Fig. S5), it was nevertheless spread across 63 contigs that cannot be scaffolded through assembly alone. Finally, these assemblies have yet to span centromeres, likely due to how they are embedded in or flanked by seas of satellite repeats (Fig. S1 in (Hoskins et al. 2015)). These limitations highlight the challenges we must overcome to resolve the most recalcitrant regions of the genome. While recent technical advances permit us to recover far more of the repetitive genome than previously, much of what we recover remains fragmented, effectively stymieing any study of structural variation. Even were we to employ the painstaking methods required to construct reasonable models of the genome structure, such progress would be incremental relative to previous results. A robust solution to these problems would entail developing approaches that can be used routinely to assemble and compare truly gapless or telomere-to-telomere genome assemblies of a population sample. The emergence of the telomere-to-telomere genome assembly and pangenome approaches (Taylor et al. 2024) brings us close to that solution.

## Methods

### DNA extraction and Sequencing

High Molecular Weight DNA was extracted from ∼200 iso-1, A4 and A3 male adult flies using Qiagen Blood and Cell Culture Midi kit following (Chakraborty et al. 2016). HMW DNA was sheared using gTUBE (Covaris) and HiFi libraries were prepared using the SMRTbell Express Template Prep Kit 2.0 (Pacific Biosciences) following the manufacturer’s recommendations. The libraries were further size selected and sequenced on the Pacific Biosciences Sequel II platform at UC Irvine Genomics High-Throughput Facility. CCS reads (ccs.bam) were generated from the Pacbio *subreads.bam* file using *ccs* (v6.2.0). CCS reads with QV>=0.99 were labeled as HiFi reads. PCR-free illumina libraries were generated from an independent DNA extraction from the strains used for HiFi reads. The libraries were sequenced on *NovaSeq 6000* platform at Novogene (Davis,CA) to generate 150 bp paired-end (2×150)reads.

### Genome Assembly

The HiFi reads longer than 6 kb were assembled using *hifiasm v0.16.1* (Cheng et al. 2021). The primary assembly from *hifiasm* was designated as our primary contig-level assembly. We removed contigs mapping to the mitochondrial genome. We aligned the first 25 kb of each contig to the *nr* database using *blastn* to identify the contigs derived from non-drosophila sources, particularly commensal microbes (Staubach et al. 2013). We then removed contigs that matched such sequences.

### Comparative and HiC based scaffolding

Two lines of evidence were employed to scaffold our iso-1 decontaminated contig assembly: reference-assisted scaffolding and HiC scaffolding. For reference-assisted scaffolding, the contig assembly was mapped to iso-1 Flybase Release 6.36 using *minimap2 v2.24* (Li 2018). The resulting alignment paf file was uploaded to *D-GENIES* web server (Cabanettes and Klopp 2018) to visualize the order and orientation of the contigs with respect to the FlyBase chromosome arm scaffolds using a dot plot. This represents our primary scaffolding approach. We also used a complementary scaffolding approach based on Hi-C contact map to incorporate sequences absent from the FlyBase reference and represented ambiguously in the dot plot. For HiC based scaffolding, we mapped the HiC reads (SRR5206663-65) from male embryos (Schauer et al. 2017) to our contig assembly and processed using the *juicer 1.6* pipeline. Subsequently *3d-dna v201008* (Dudchenko et al. 2017) was used to perform scaffolding. Additionally, the HiC contact map was compared to the D-GENIES dot plot to resolve ordering discrepancies, with reference-guided ordering given priority over Hi-C. The information from both synteny and HiC were integrated to generate the final scaffolded reference using a custom script. The final HiC contact map for iso-1 can be found in Supp Fig. 4. A4 and A3 decontaminated contig assemblies were mapped to iso-1 scaffolded HiFi assembly using *minimap2 v2.24*. Reference assisted scaffolding was performed using the procedure described above.

### Quality Control and Heterozygosity Analysis

We calculated BUSCO and consensus quality (QV) to test the quality of our assembled genomes. The BUSCO scores for the decontaminated contig assemblies were calculated using *compleasm v0.2.2* (Huang and Li 2023) (Supp Table 5). *Merqury v1.3* (Rhie et al. 2020) was used to obtain phred QV score and k-mer completeness for the final scaffolded assemblies (Supp Table 5). The Illumina data generated for the 3 strains was used during *Merqury* evaluation. The QV score reported here is the average of the major chromosomal arms. Heterozygosity analysis was performed using *GenomeScope 2.0* (Ranallo-Benavidez et al. 2020). We tweaked the standard approach to derive heterozygosity estimates for our datasets. Since we had sequenced the heterogametic sex (males), *GenomeScope* interpreted the haploid peak (from X and Y chromosomes) as the heterozygous peak. Thus, we employed only reads derived from autosomes. HiFi reads were mapped to the scaffolded genome using *minimap2 v2.25* and reads mapping to autosomes were extracted. This subset of reads were further used to count kmers (using *Jellyfish v2.3.0* (Marçais and Kingsford 2011)) and obtain a *GenomeScope* profile (Supp Fig 1,2,3).

### Identification of Y-linked contigs

We employed a modified approach from Chang & Larracuente (2019) to identify Y-linked contigs. Male and Female PCR-free Illumina reads (SRR6399448-49) were mapped to the contig assemblies using *bwa mem 0.17.7*. *Samtools coverage 1.16 (-q 20)* was used to calculate coverage statistics for each contig in the assembly. Among the various metrics reported by *samtools coverage* (e.g., *covbases, coverage, meandepth*), we used the *coverage* metric as it provides the percentage of the contig covered by reads. We used the ratio of male to female *coverage* to determine the sex-linkage of the contigs. For autosomal contigs and X this ratio should be ∼1, but for Y linked contigs the ratio should be significantly greater than 1. We classified any contig with *coverage* ratio > 2 as Y-linked (excluding contigs with less than 5 reads mapped). Contigs in females that have 0 *coverage*, but have *coverage* > 0 in males were also classified as Y-linked. For contigs (such as consisting entirely of simple sequence repeats), which have 0 *coverage* in both male and female when a filter of MAPQ > 20 is applied, we applied the procedure with no MAPQ cutoff. Such contigs with 0 *coverage* in females and non-zero coverage in males were classified as Y-linked.

### Synteny, Repeat Content and Depth Analysis

Synteny analysis was performed using *D-Genies* (Cabanettes and Klopp 2018). The genomes were aligned using *minimap2 v2.24* to obtain a *paf* file. Index files for the genomes were created by a python script provided by the *D-Genies* authors. The resultant paf file and index files were uploaded to the *D-Genies* server (Anon). The online server is an interactive way to visualize synteny using a dot plot and generate static dot plot images.

*RepeatMasker v4.1.2-p1* (Smit et al. 2015) was used to identify repeats in our assemblies. The default Dfam 3.3 curated library (Storer et al. 2021) was augmented with a few sequences relevant to our study, such as Stellate, Histone, rRNA, to obtain a custom repeat library. This custom library was used to identify repeats in our assemblies. The extra sequences appended can be found on the GitHub page.

We used depth analysis to verify the integrity of an assembled sequence. HiFi reads were mapped to the assembly using *minimap2 v2.24*, and *samtools depth v1.6 (Danecek et al. 2021)* was used to calculate per-base depth for the locus. An Rscript was used to plot and visualize the depth across the locus.

### Targeted Assembly of the Histone cluster

HiFi reads were mapped to the entire contig assembly using *minimap2 v2.24*. The list of reads mapping to the histone locus +10 kb flank was obtained using *samtools view*. *seqtk v1.3-r116-dirty* was used to extract the sequences in the list file. These reads were assembled using *hifiasm 0.16.1* with repeat sensitive and targeted assembly parameters (*-D 10 –hg-size <estimated_size_locus>*). The largest contig from the *primary assembly (.p_ctg.fasta)* output was considered as our histone locus. In addition, *the unitig graph (*.p_utg.gfa*) from both the default and targeted assemblies was visualized in *Bandage v0.9.0* in order to investigate the structure of the locus further (Supp Fig 9,12,14). For A3, we also assembled the longest 40X reads but both approaches failed to assemble the entire cluster. Thus, we decided only to consider the well-resolved parts of the 40X *unitig graph* for our final assembly (Supp Fig 14). The nodes in the path (shown by red lines) were merged in *Bandage v0.9.0,* and their fasta sequences were written to the disk. The sequences were finally scaffolded together in proper order and orientation using our scaffolding method.

### Inferring histone copy number from short read data

The Histone Locus in iso-1 HiFi scaffolded assembly was masked. A single histone unit (5kb type) was added as a separate contig to the assembly. The short read Illumina data was mapped to the modified reference using *bwa mem v0.7.17 (Li 2013)*. *Samtools depth v1.6 (-aa -J -s)* was used to calculate the per base depth for the Histone contig and the 2L arm. The ratio (Histone_contig/2L) of the median per base depth was inferred to be the histone copy number for that particular strain. The SRR IDs of the GDL strains used here are provided in Supp File 2.

### Analysis of histone locus using phylogenetic methods

The beginning and start sequences of a single Histone Unit were identified. *RepeatMasker* was used to find their locations in all the histone arrays using these sequences. The individual units from each histone loci were extracted using the RepeatMasker output file. The TE-containing units were removed, and all units from the two strains were merged into a single fasta file for pairwise comparison. The combined fasta file was imported into *MEGA v11.0.13* (Tamura et al. 2021), and a multiple sequence alignment (MSA) was performed using *MUSCLE*. This MSA was used as input for building a phylogenetic tree. A Neighbor-Joining tree (500 bootstraps) was constructed using default parameters. Finally, a 50% bootstrap consensus tree was obtained. If only two units, one from each strain, clustered together in the consensus tree, they were marked as putative anchors. This procedure was followed for all pairwise comparisons. The blue boxes highlight such pairs in the pairwise phylogenetic trees (Supp File 3). To identify closely related repeat units (Fig 4B), we looked at clades containing units from both strains in each pairwise comparison. We correlated results across all the pairwise comparisons to obtain an overall pattern for the 3 strains. The red, purple, and green boxes highlight such clades (Supp File 3). The colors here correspond to the ones used in Fig 4B. The molecular distances used to plot Fig 4C were obtained using MEGA using default parameters.

### Linkage Disequilibrium analysis in and around histone locus

Illumina short reads (Supp File 2) were trimmed using *TrimGalore v0.6.10* (Krueger). The trimmed reads were mapped to the iso-1 HiFi scaffolded assembly, with its Histone locus masked, using *bwa mem v0.7.17.* Variants were called using *octopus v0.7.4 (Cooke et al. 2021)* for a ∼1 Mb region, flanking the histone cluster for each individual strain. Joint genotyping was performed using previously generated *bcf* files for 147 strains. A few filters were applied to the jointly called vcf using *bcftools v1.7* (Danecek et al. 2021). Genotypes that were not labeled ‘PASS’ were converted to ‘no calls’; only biallelic sites were further considered; heterozygous calls were converted to ‘no calls’ since we are dealing with haploid embryos. Finally, sites with missing genotype calls greater than 5% or having minor allele frequency less than 0.05 were filtered out. This vcf file was imported into *IGV* (Thorvaldsdóttir et al. 2013) for further analysis. We used *PLINK v1.90b7* (Chang et al. 2015) with biallelic SNPs to calculate the pairwise LD (r^2^) matrix.

### Analysis of stellate locus using phylogenetic methods

The same methods used to analyze the histone locus were applied to investigate the stellate locus. The blue boxes represent the anchors identified for the euchromatic and heterochromatic locus for iso-1 HiFi, A4 HiFi and A3 HiFi (Supp File 4). A similar procedure was followed for comparing the euchromatic and heterochromatic stellate locus from 3 iso-1 assemblies (Supp File 5).

## Data Access and Code Availability

All raw reads and genome assemblies are deposited to NCBI under Bioproject PRJNA1122219. All scripts, analysis pipelines, and relevant data files are deposited in GitHub (https://github.com/harsh-shukla/Dmel_HiFi_Asm_variation).

## Acknowledgments

We thank Kevin Thornton, Grace YC Lee, and Yuheng Huang for the discussions. We also thank Yuheng Yuhang, Chris Fiscus, Doc Edge, Matt Pennell, and Ching-Ho Chang for their comments on an early version of the manuscript. This work was funded by the National Institutes of Health (R00GM129411 to M.C.; R01GM123303 and R35GM153327 to J.J.E.) and start-up funding from Texas A&M to M.C. This work utilized resources of the UCI Genomics Research and Technology Hub (GRT Hub) parts of which are supported by NIH grants to the Comprehensive Cancer Center (P30CA-062203) and the UCI Skin Biology Resource Based Center (P30AR075047) at the University of California, Irvine, as well as to the GRT Hub for instrumentation (1S10OD010794-01and 1S10OD021718-01).

