## Supplementary material for "Genetic variation in recalcitrant repetitive regions of the *Drosophila melanogaster* genome": Supp File 5

### ISO1 HiFi and Rel6 Euchromatin Stellate locus

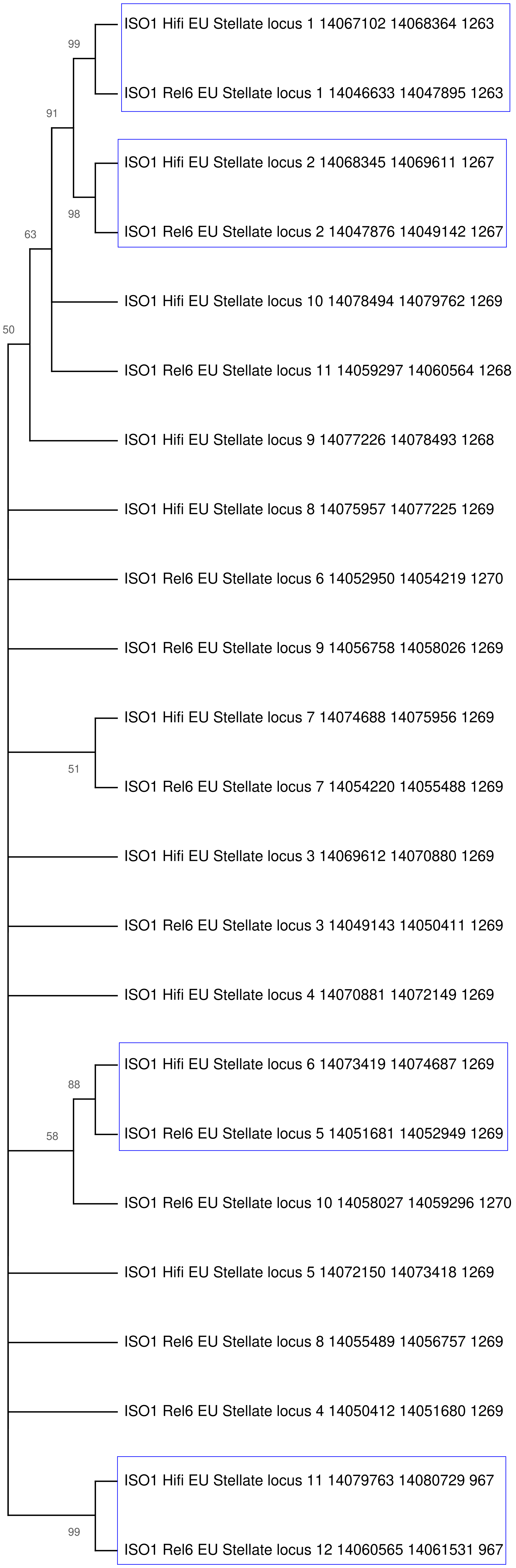

### ISO1 HiFi and HetEnr Euchromatin Stellate locus

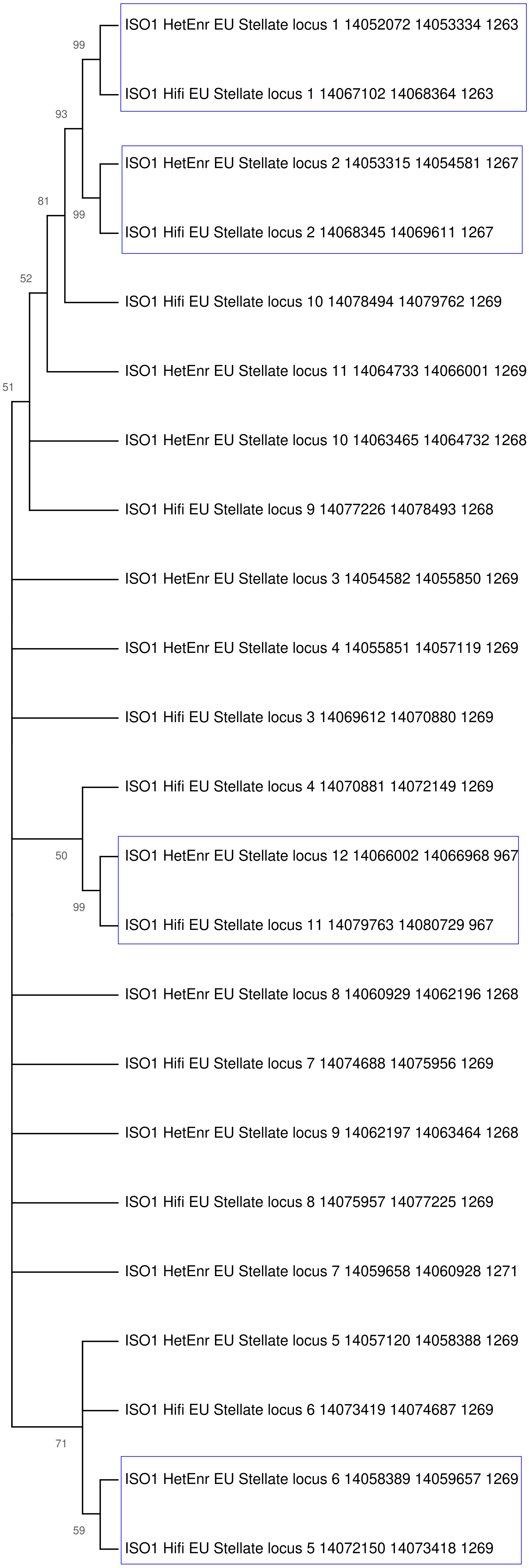

### ISO1 HiFi and HetEnr Euchromatin Stellate locus

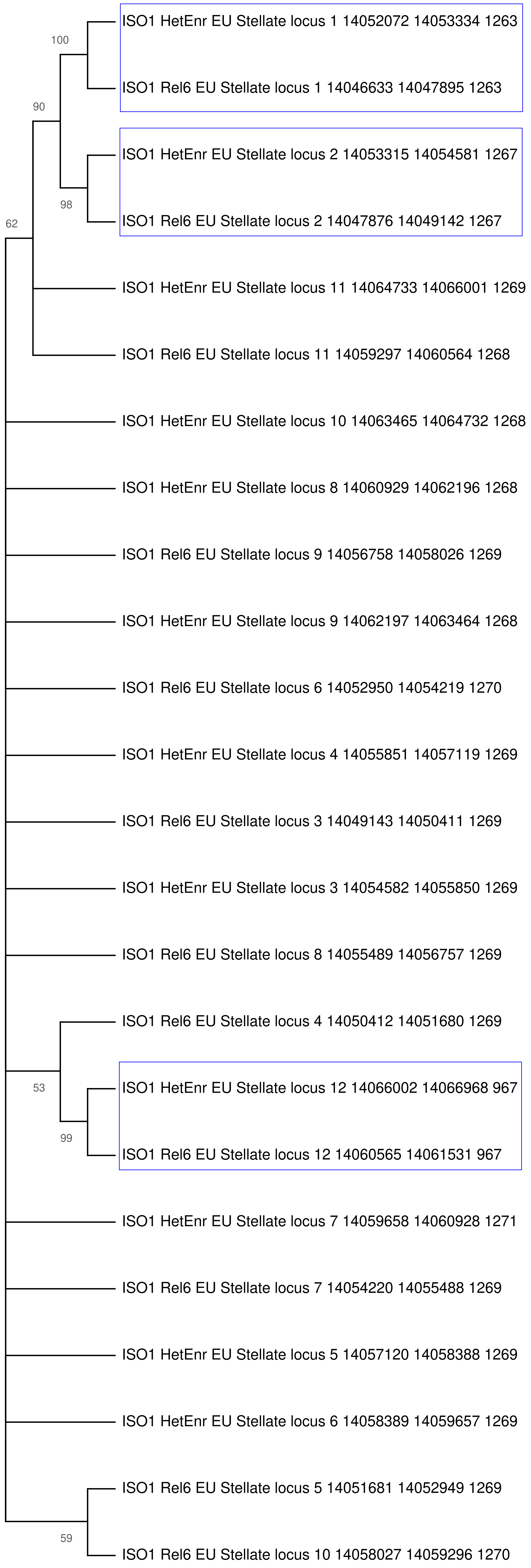

### ISO1 HiFi and HetEnr Het. L2 Stellate locus

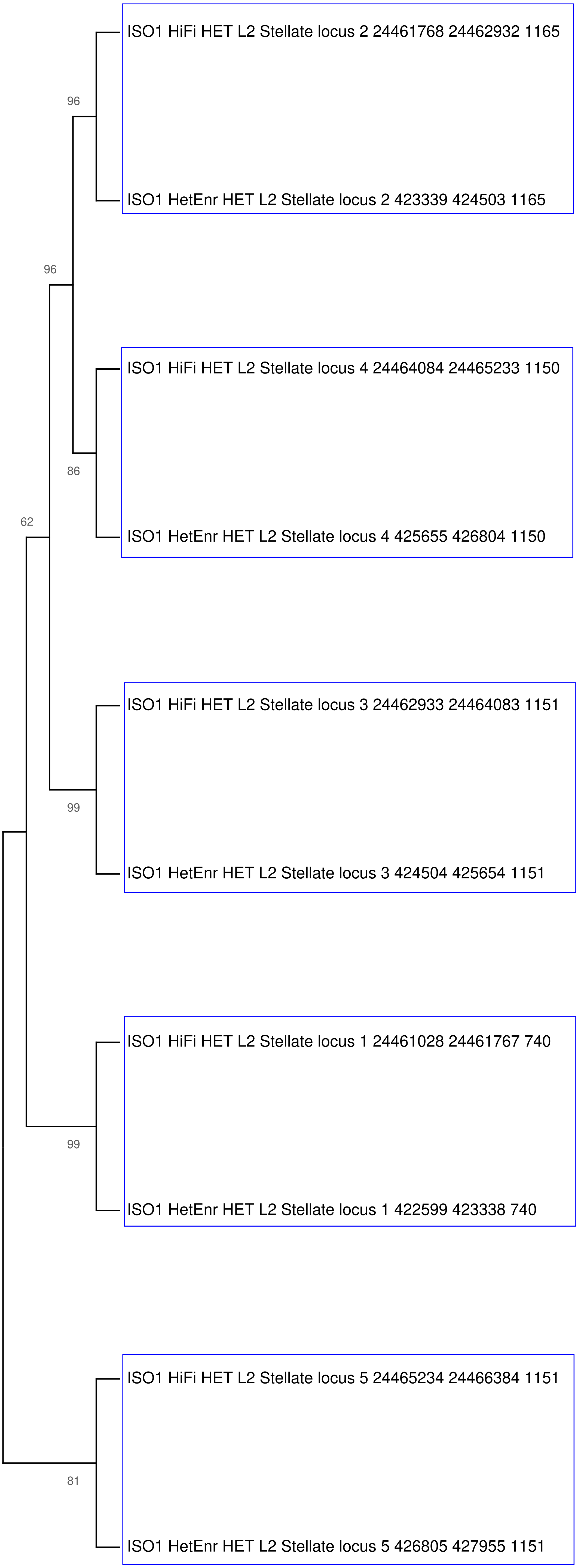

### ISO1 HiFi and HetEnr Het. L3 Stellate locus

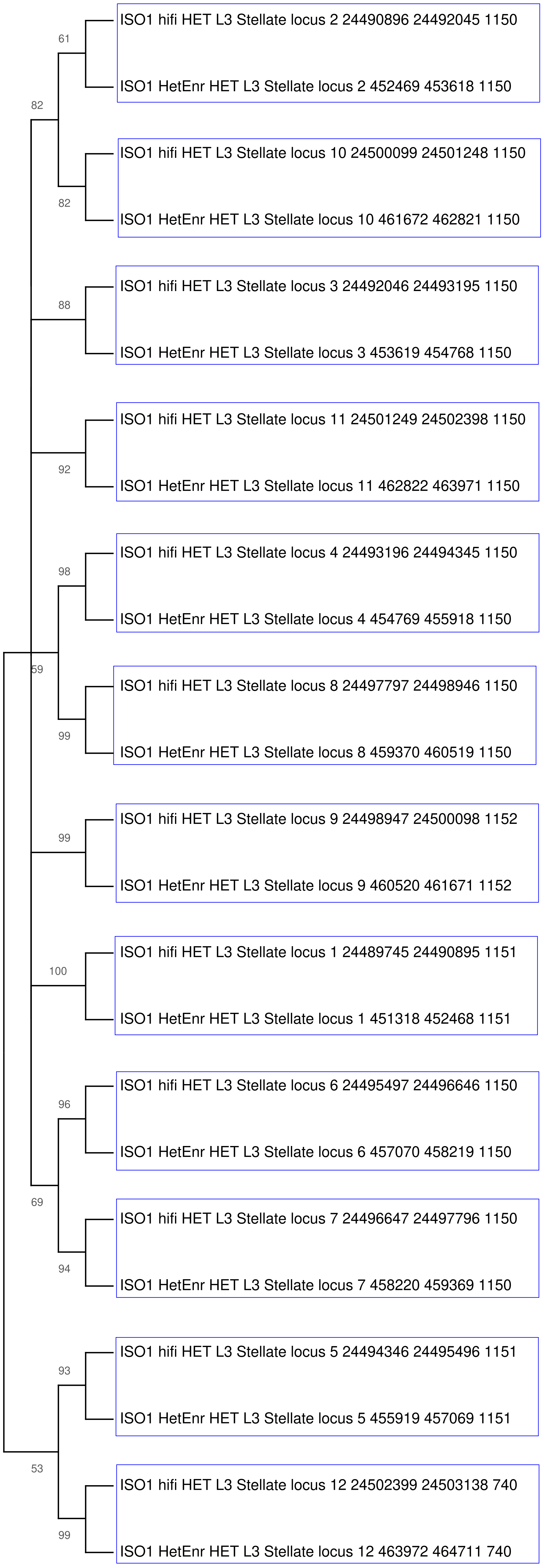
