## Supplementary figures and images for "Genetic variation in recalcitrant repetitive regions of the *Drosophila melanogaster* genome"

### Supp File 3

#### ISO1 HiFI - ISO1 Rel6

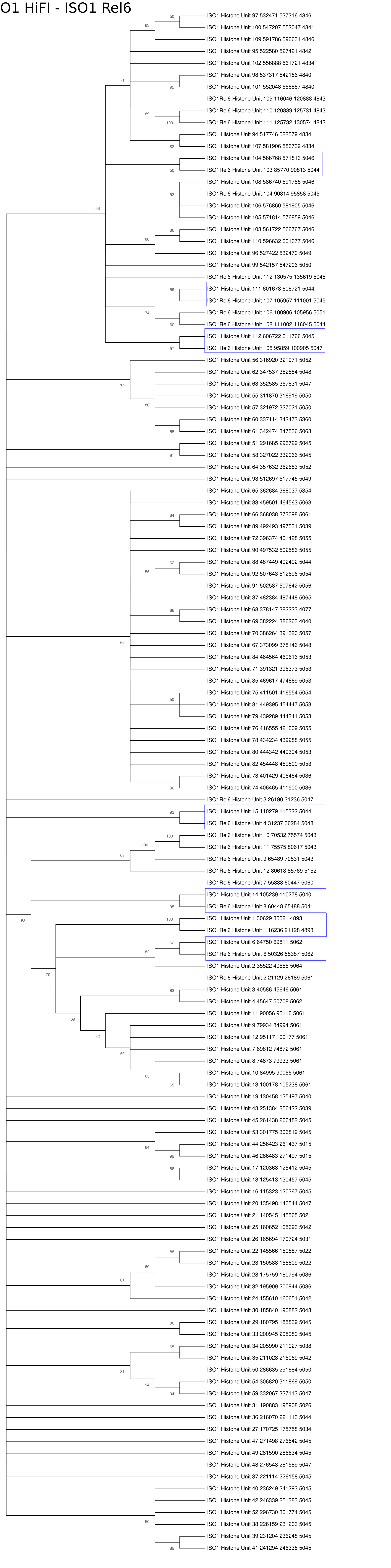

#### ISO1 HiFi - A4 HiFi

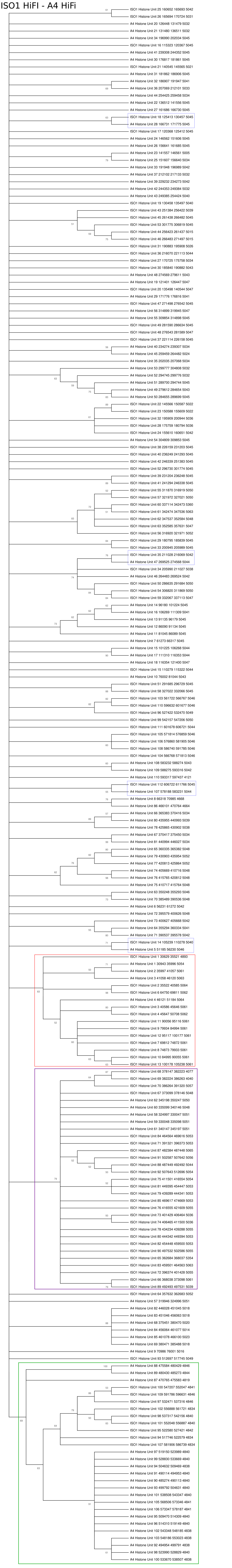

#### ISO1 HiFi - A3 HiFi

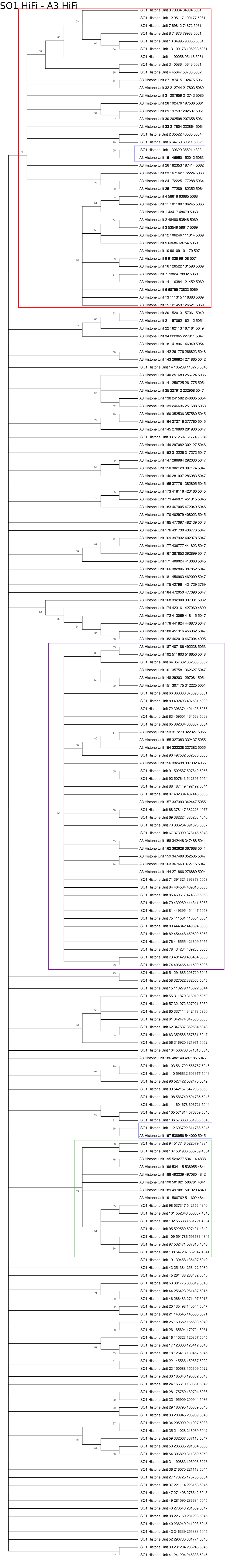

#### A4 HiFi - A3 HiFi

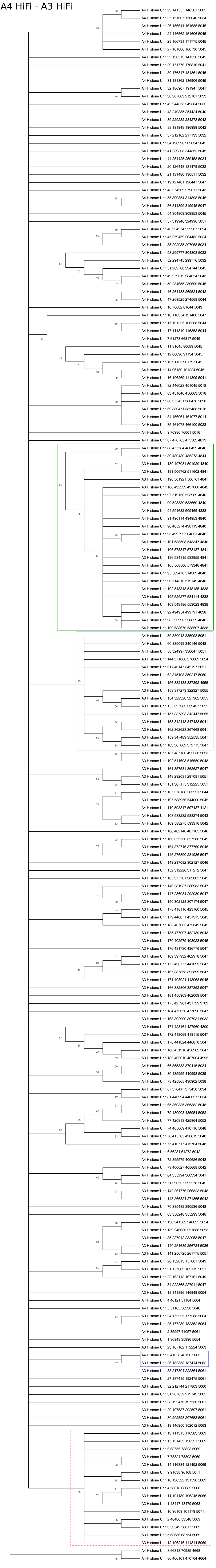

### Supp File 4

#### ISO1 A4 A3 Euchromatic Stellate Locus

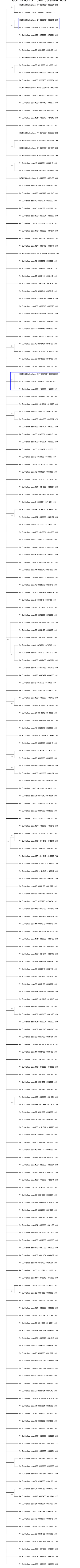

#### ISO1 A3 Het. L2 Stellate Locus

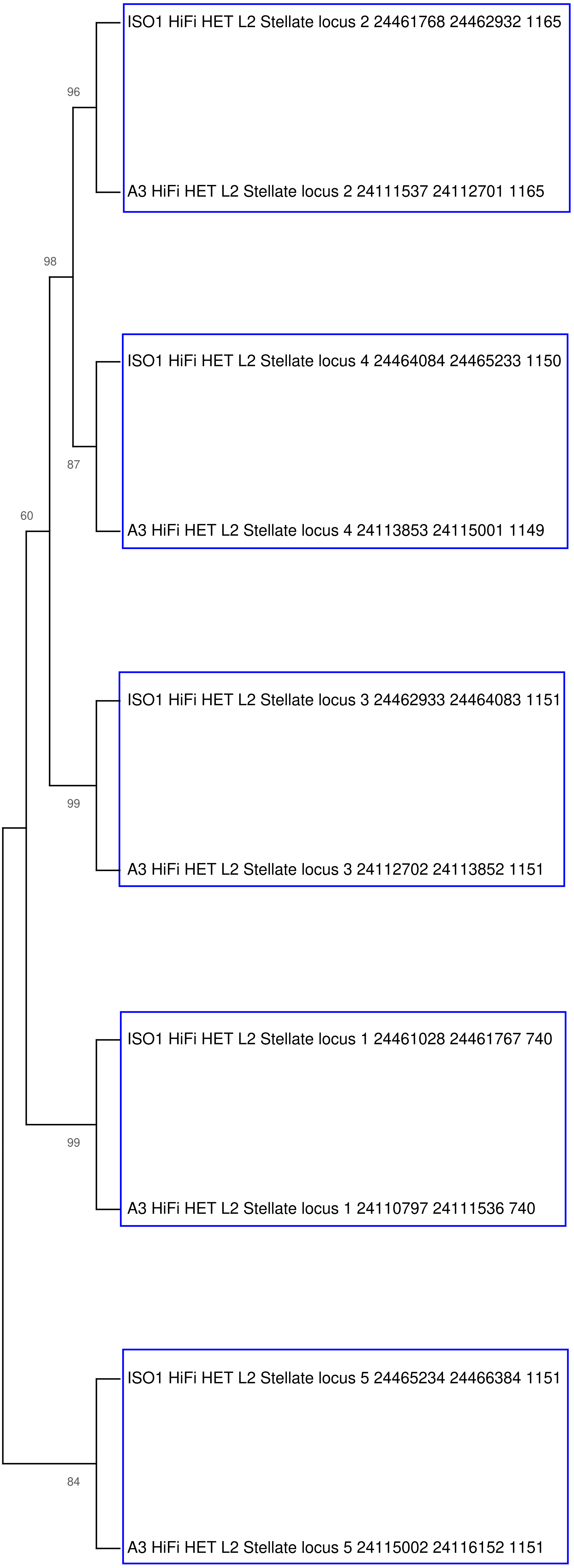

#### ISO1 A3 Het. L3 Stellate Locus

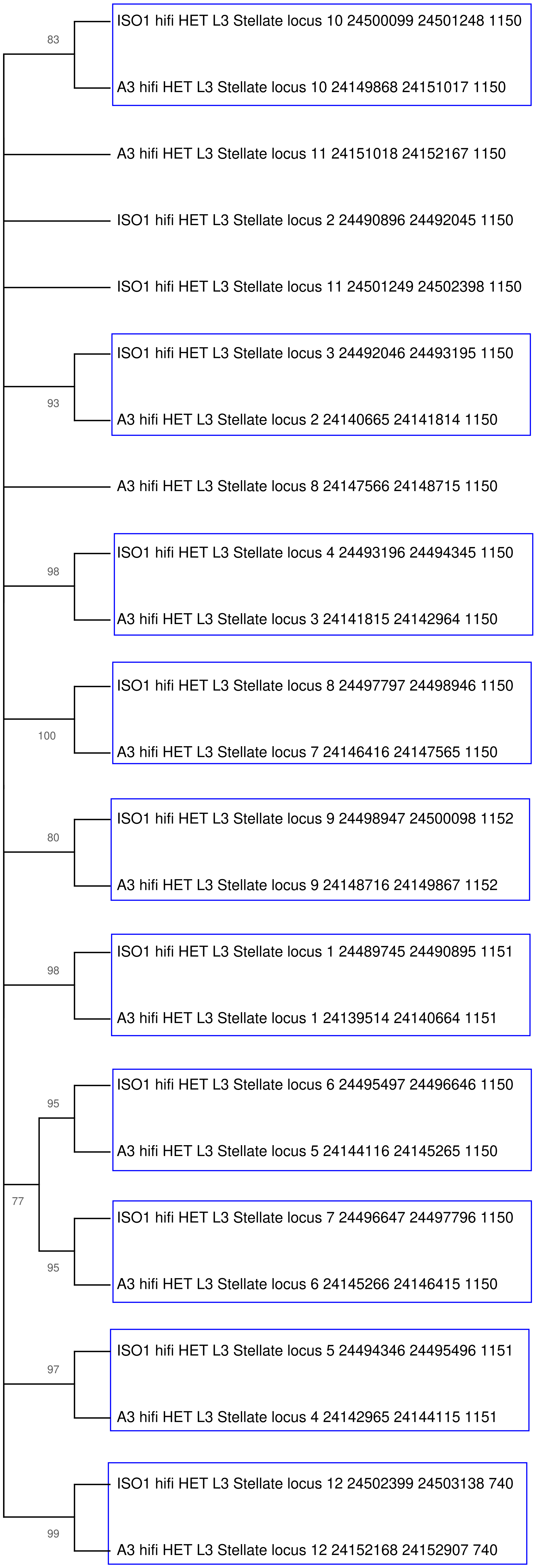
