## Supplementary material for "Genetic variation in recalcitrant repetitive regions of the *Drosophila melanogaster* genome": Supp Fig

| Strain | Throughput (Gb) | Median | Q1 | Q2 | Approx. Depth |
|---|---|---|---|---|---|
| iso-1 | 12.40 | 15,026 | 13,250 | 17,363 | 97x |
| A4 | 21.28 | 13,724 | 12,013 | 15,779 | 163x |
| A3 | 20.75 | 15,679 | 12,920 | 19,663 | 156x |

**Supplementary Table 1.** The statistics of HiFi datasets generated in this study

| Strain | Assembly size (Mb) | Num of contigs | N50 (Mb) | N90 (Mb) | L50 | L90 | Largest (Mb) |
|---|---|---|---|---|---|---|---|
| iso-1 | 168.56 | 236 | 22.96 | 0.463 | 4 | 33 | 27.23 |
| A4 | 170.31 | 277 | 21.53 | 0.336 | 4 | 38 | 28.16 |
| A3 | 172.45 | 142 | 21.50 | 0.831 | 4 | 20 | 28.05 |

**Supplementary Table 2.** The statistics of contig level assemblies (after decontamination and removal of redundant mitochondrial contigs)

| Strain | Assembly size (Mb) | No. of Scaffolds | N50 (Mb) | N90 (Mb) | L50 | L90 | Largest (Mb) | No. of Ns | No. of Gaps |
|---|---|---|---|---|---|---|---|---|---|
| iso-1 | 168.58 | 212 | 27.44 | 0.697 | 3 | 13 | 34.07 | 24,000 | 24 |
| A4 | 170.33 | 259 | 27.72 | 0.368 | 3 | 21 | 34.41 | 18,000 | 18 |
| A3 | 172.46 | 126 | 28.05 | 1.33 | 3 | 11 | 34.39 | 16,000 | 16 |

**Supplementary Table 3.** The statistics of Scaffolded assemblies

| CHR | iso-1 Rel6 | iso-1 HiFi | A4 HiFi | A3 HiFi |
|---|---|---|---|---|
| 2L | 23,513,712 | 24,266,334 | 24,085,515 | 23,885,995 |
| 2R | 25,286,936 | 25,878,355 | 25,269,289 | 25,586,733 |
| 3L | 28,110,227 | 28,948,966 | 28,897,970 | 28,636,850 |
| 3R | 32,079,331 | 34,070,894 | 34,418,861 | 34,392,019 |
| 4 | 1,348,131 | 1,303,249 | 1,416,745 | 1,393,402 |
| X | 23,542,271 | 27,447,821 | 27,727,683 | 28,052,005 |
| Y | 3,667,352 | 15,707,245 | 16,486,182 | 17,032,472 |

**Supplementary Table 4.** The length of major chromosome of Scaffolded HiFi assemblies compared to iso-1 Release 6

| Assembly | Complete - Single Copy | Complete - Duplicated | Fragmented | Missing | QV (arms) | Kmer-completeness |
|---|---|---|---|---|---|---|
| iso-1 Rel6 | 3265 (99.39%) | 10 (0.30%) | 8 (0.24%) | 2 (0.06%) | - | - |
| iso-1 HiFi | 3266 (99.42%) | 9 (0.27%) | 8 (0.24%) | 2 (0.06%) | 49.9 | 99.2574 % |
| A4 HiFi | 3261 (99.27%) | 14 (0.43%) | 8 (0.24%) | 2 (0.06%) | 56.1 | 99.3543 % |
| A3 HiFi | 3266 (99.42%) | 9 (0.27%) | 8 (0.24%) | 2 (0.06%) | 55.3 | 99.33 % |

**Supplementary Table 5.** The BUSCO score of the HiFi assemblies and iso-1 Release 6 (from *compleasm*) . Phred QV scores and kmer-completeness for the hifi assemblies. The QV score is average QV of major chromosomal arms (2L, 2R, 3L, 3R, X)

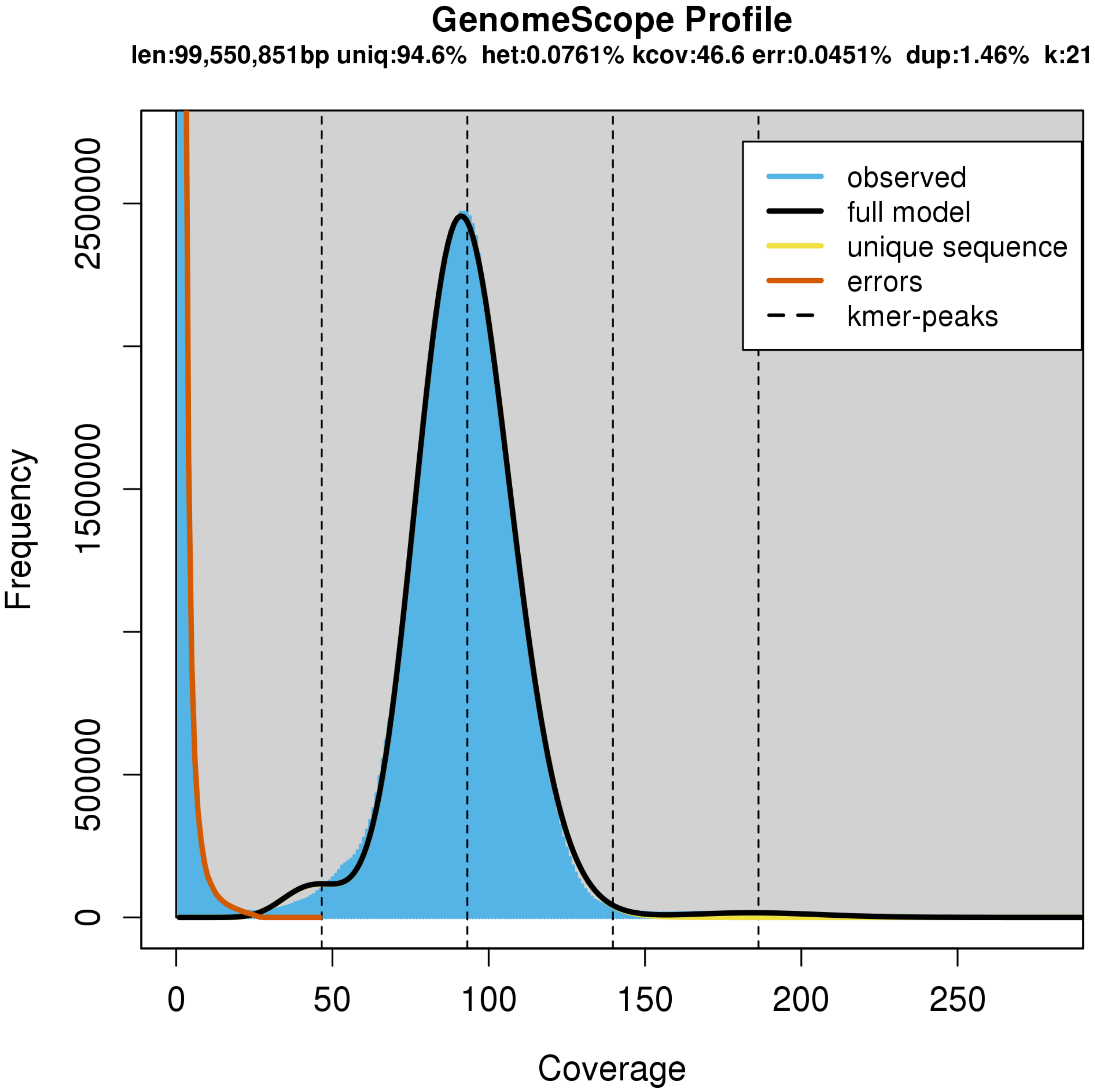


**Supplementary Fig 1.** Estimation of heterozygosity using genomescope - iso-1

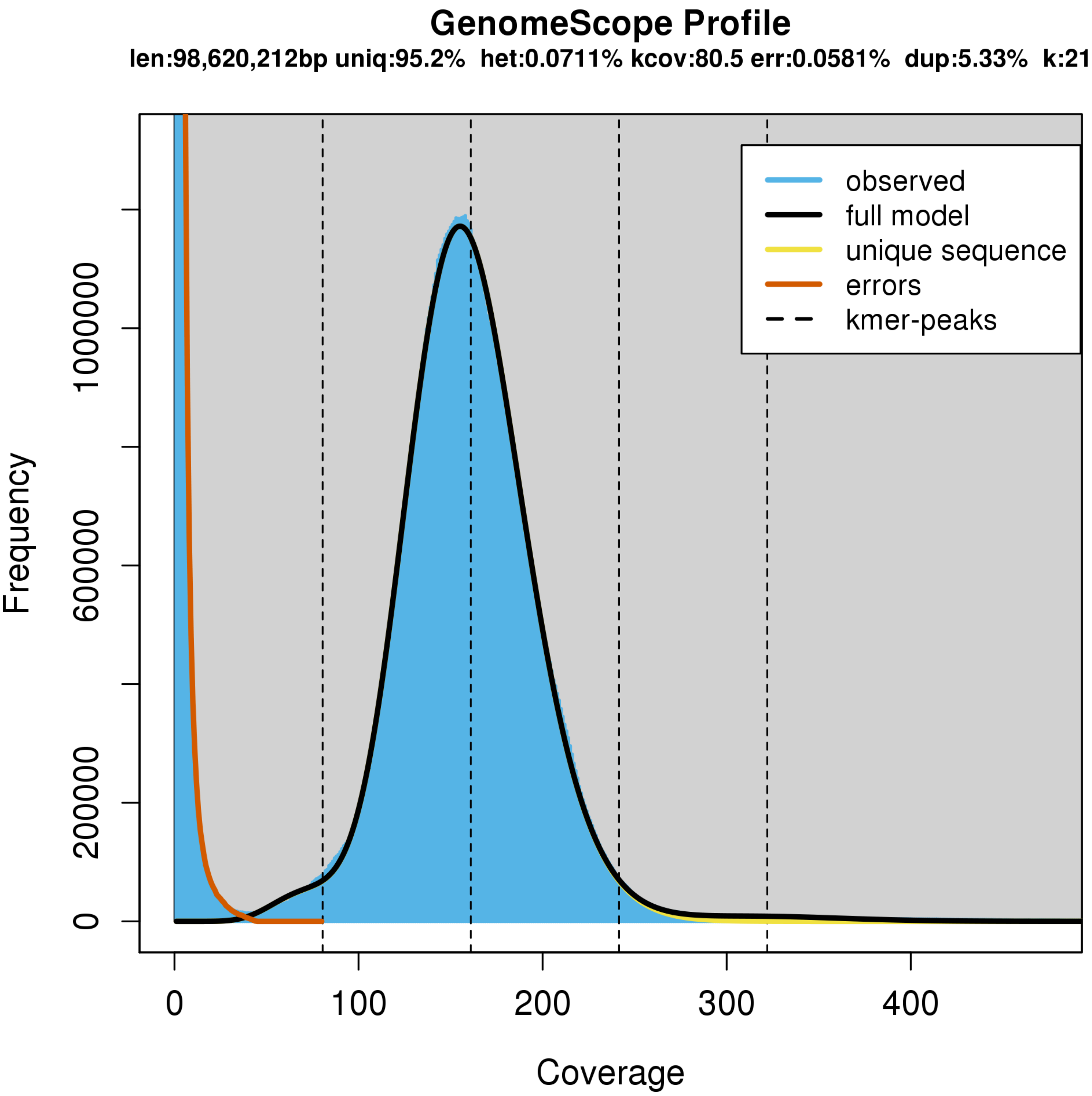


**Supplementary Fig 2.** Estimation of heterozygosity using genomescope - A4

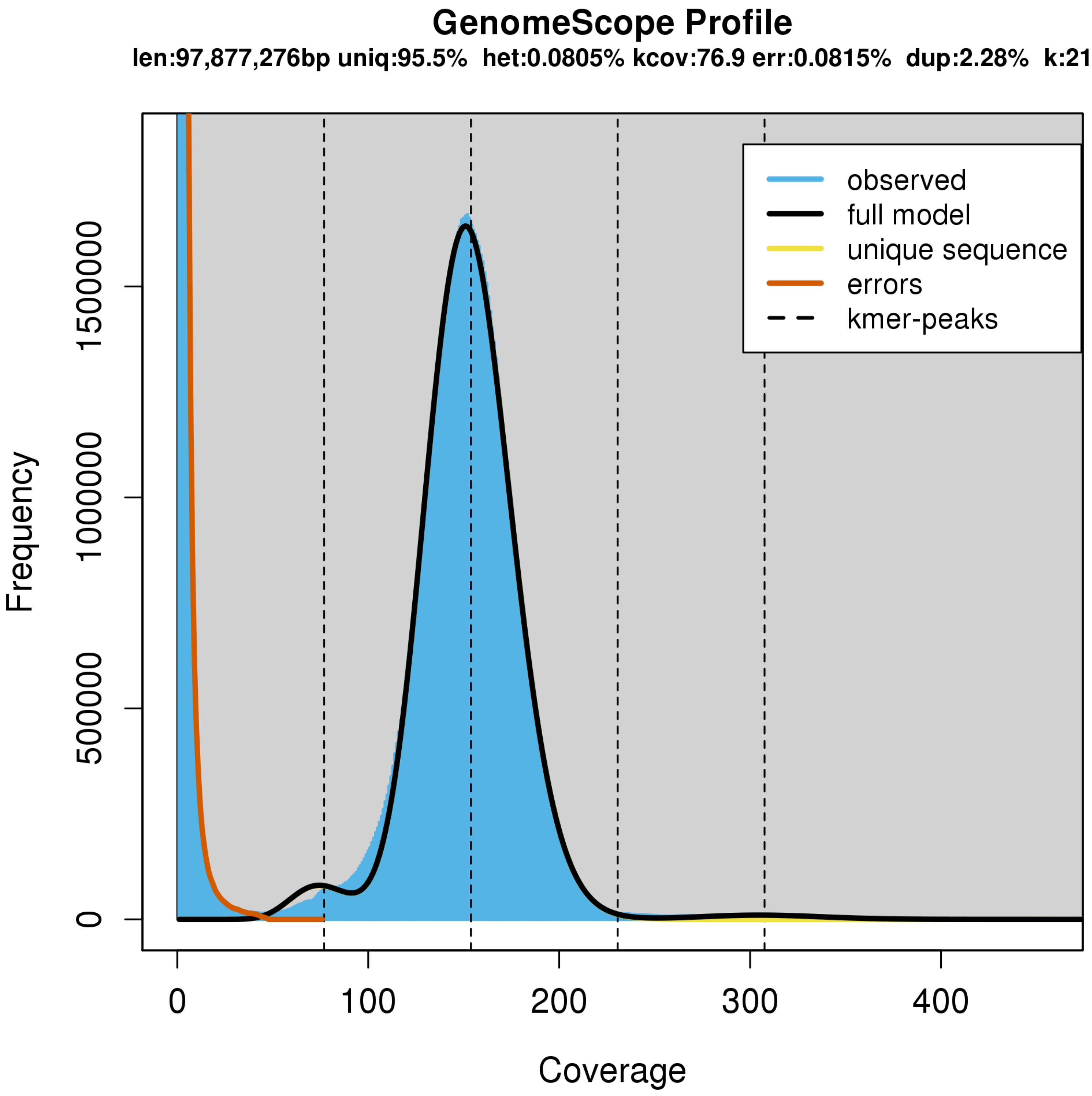


**Supplementary Fig 3.** Estimation of heterozygosity using genomescope - A3

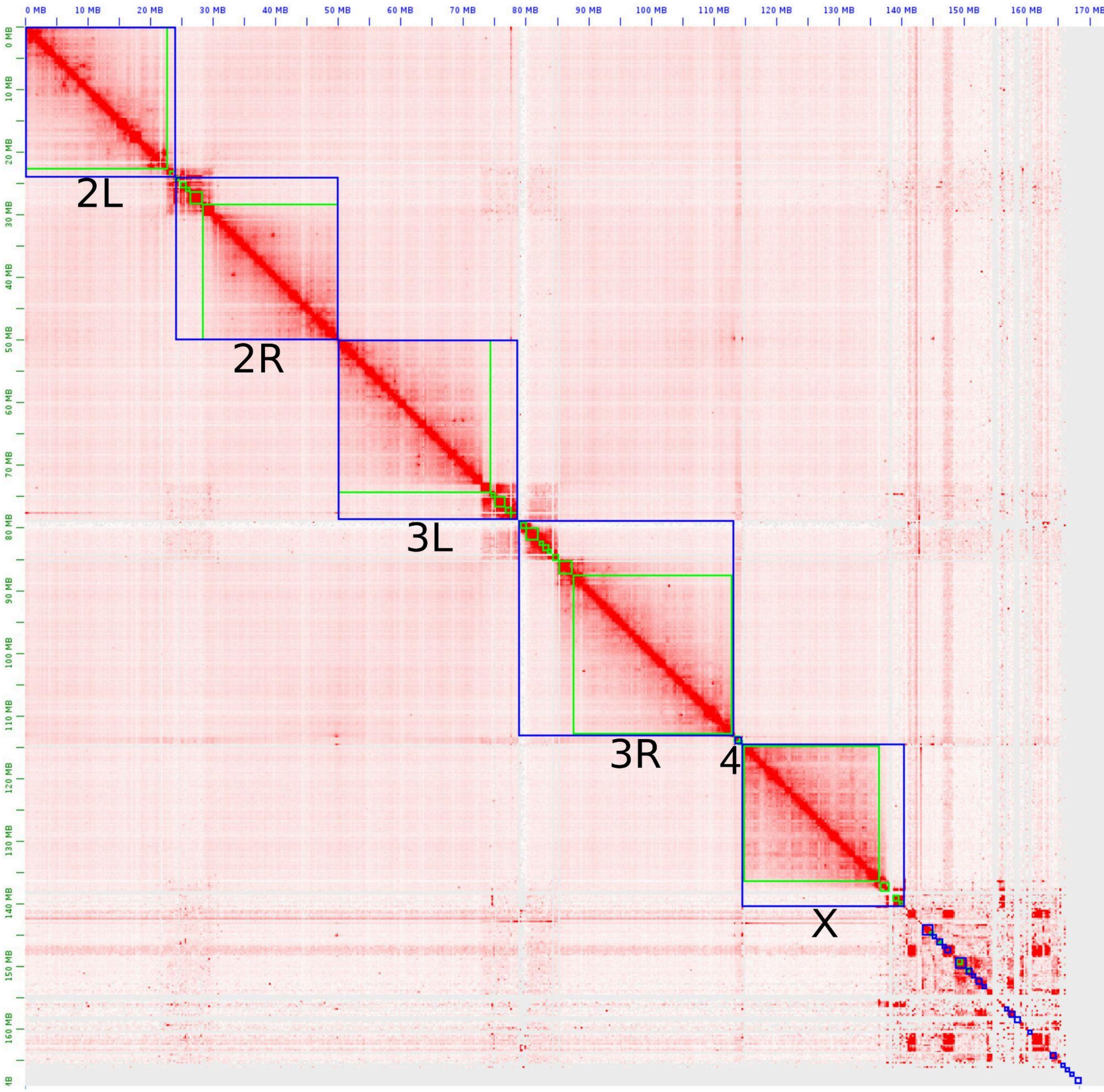


**Supplementary Fig 4.** HiC Scaffolding of iso-1 hifi contig assembly

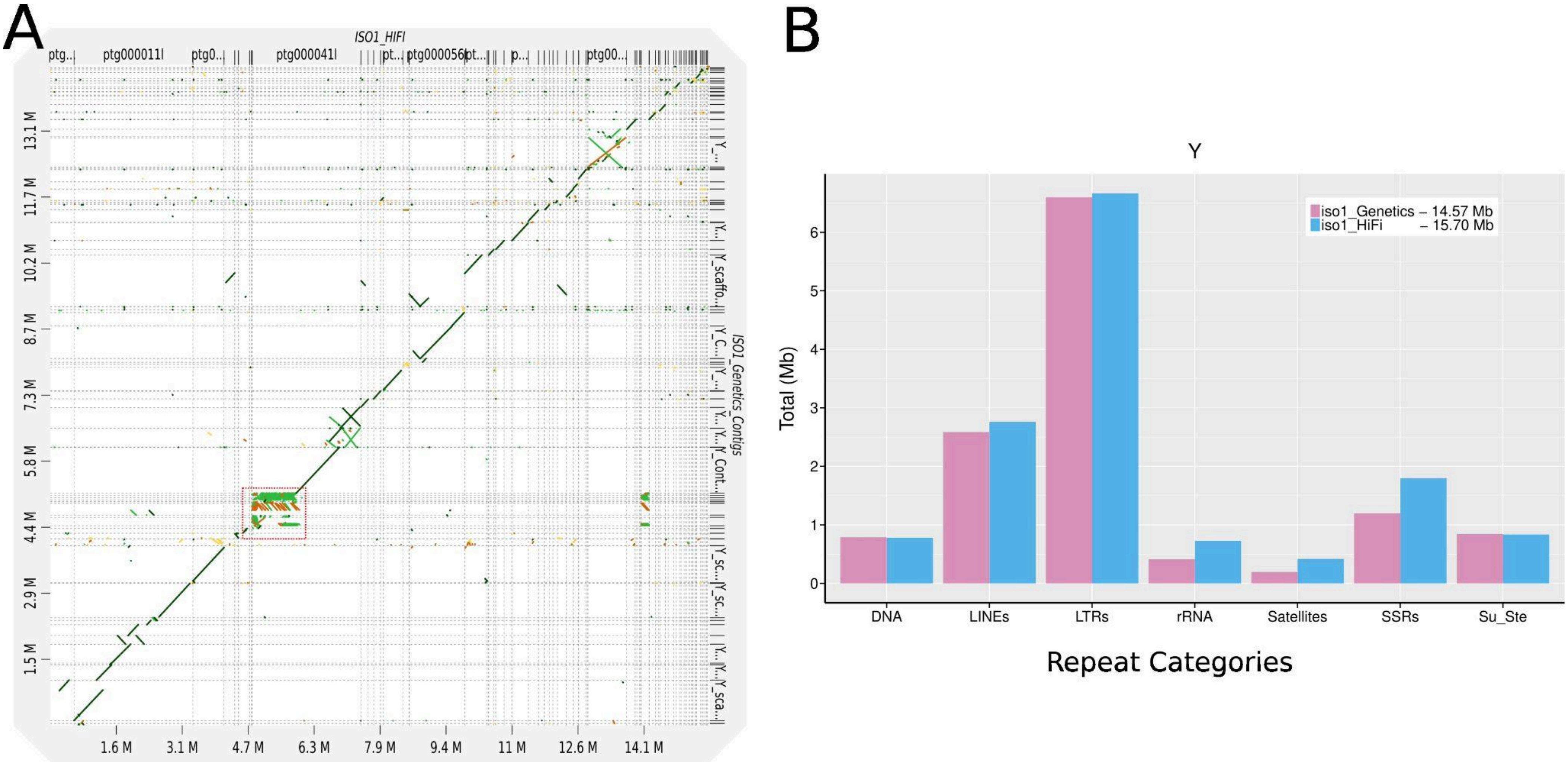


**Supplementary Fig 5A.** Genomic dot plot showing synteny between iso-1 HiFi Y-contigs (X-axis) vs iso-1 Genetics Y-contigs (Y-axis).

**Supplementary Fig 5B.** Repeat Content comparison between the two Y assemblies. They are very similar. iso-1 Hifi assembly just has more rRNA sequences assembled and as a consequence has more rRNA, rRNA associated satellites and rRNA associated R1,R2 LINEs. The beginning of contig 041l (red box) is the rRNA assembled locus. ~80% of the Y chromosome consists of simple tandem repeats. Our iso-1 HiFi assembly also recovers more tandem repeats (SSRs).

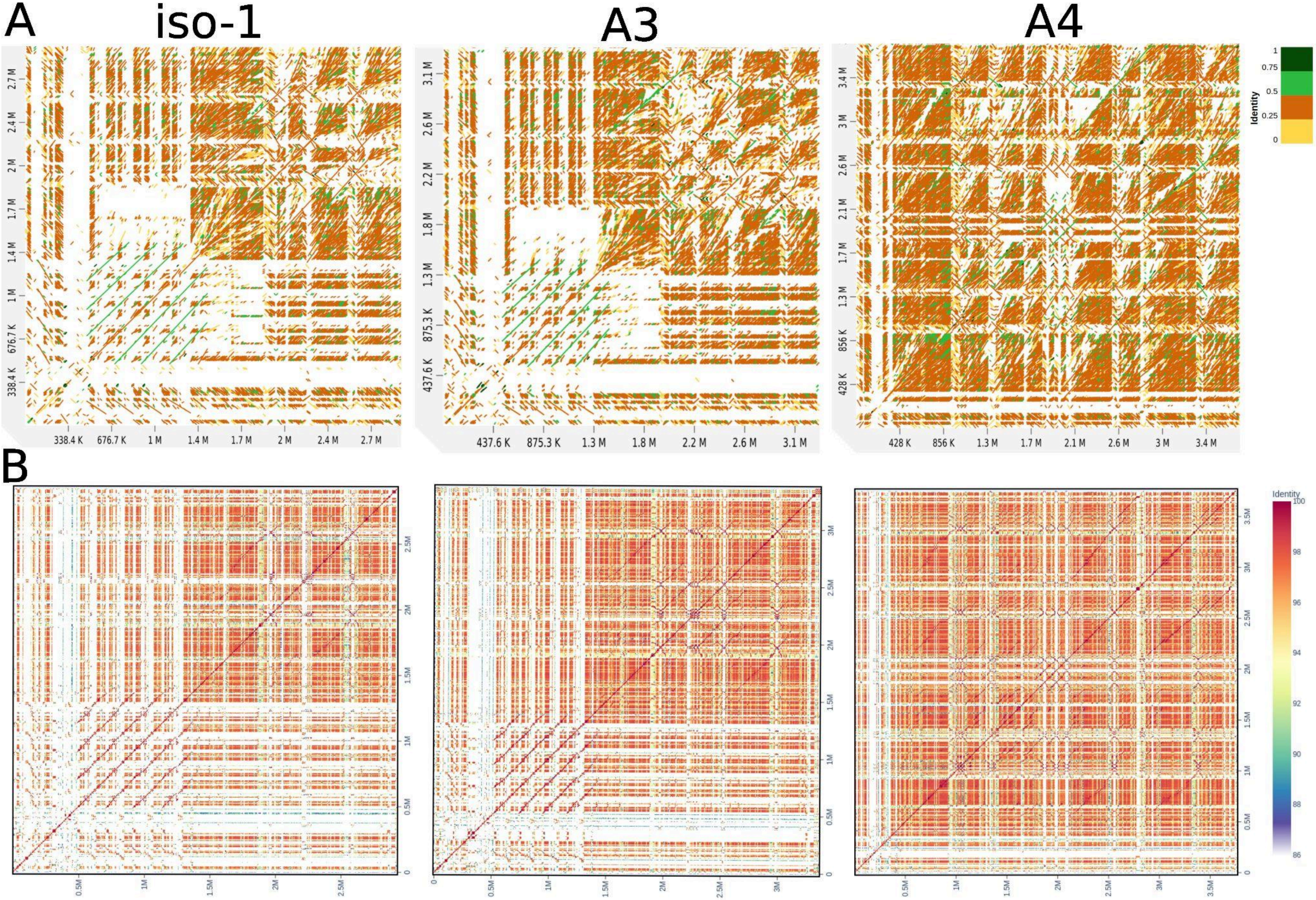


**Supplementary Fig 6 A and B.** Intra-strain self dot plots of newly assembled X sequences (excluding the rDNA array) in the three assembled strains. The X and Y axis for each plot are identical (the newly assembled X sequence for the corresponding strain). Plots in **A** are from *D-Genies* and plots in **B** are from *ModDotPlot.*

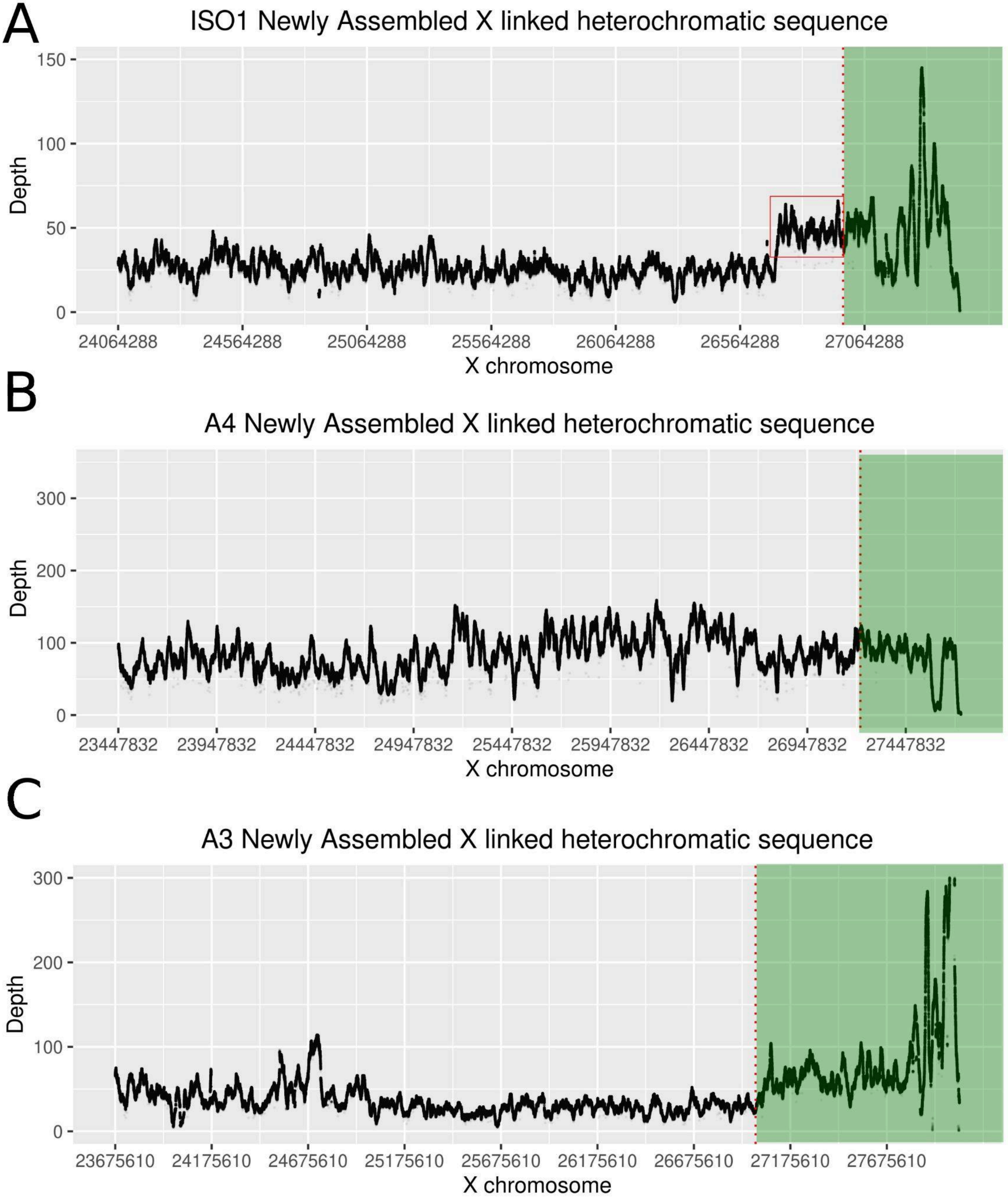


**Supplementary Fig 7.** The depth plots(using HiFi reads) of newly assembled X linked sequences in the 3 strains. The green box represents the rDNA cluster. The red box in iso-1 hints at a possible ~250 kb collapse in our iso-1 hifi assembly

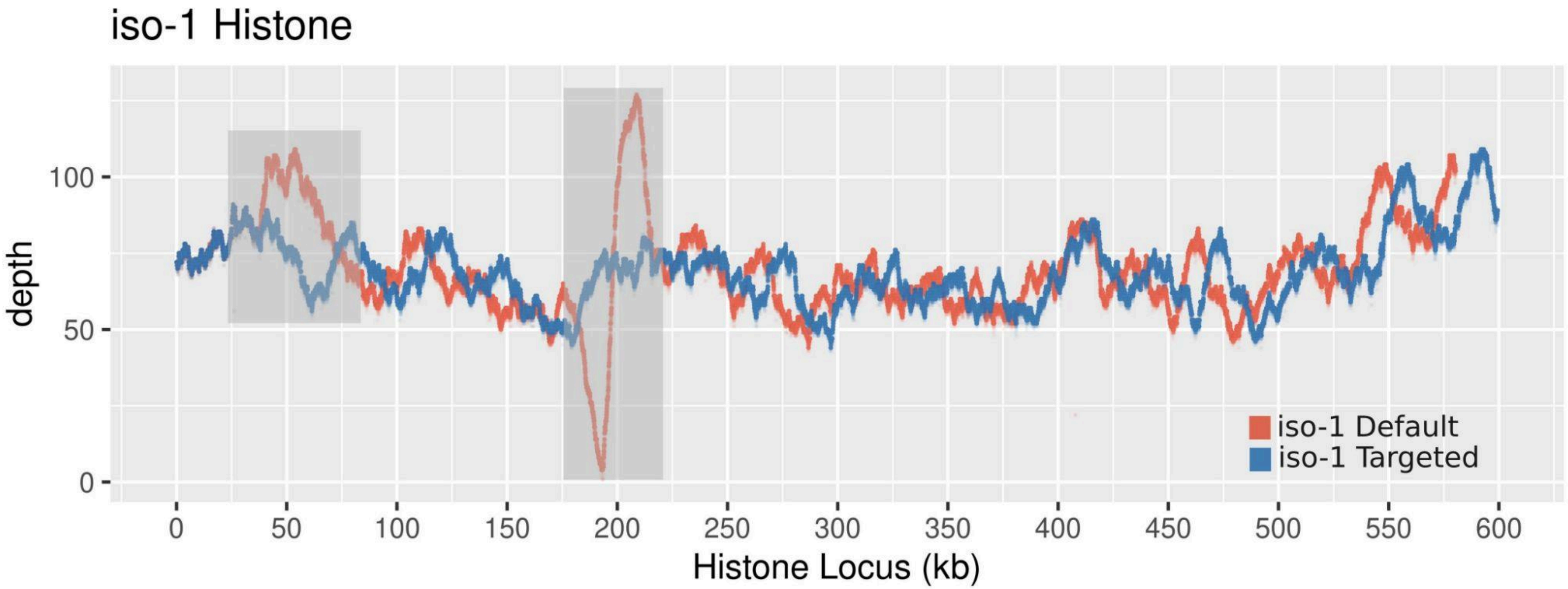


**Supplementary Fig 8.** The depth plots of HiFi reads mapped to the Default and Targeted histone locus for iso-1. The gray boxes highlight a possible collapse in the beginning of the cluster (~50kb) and a misassembly (~200kb).

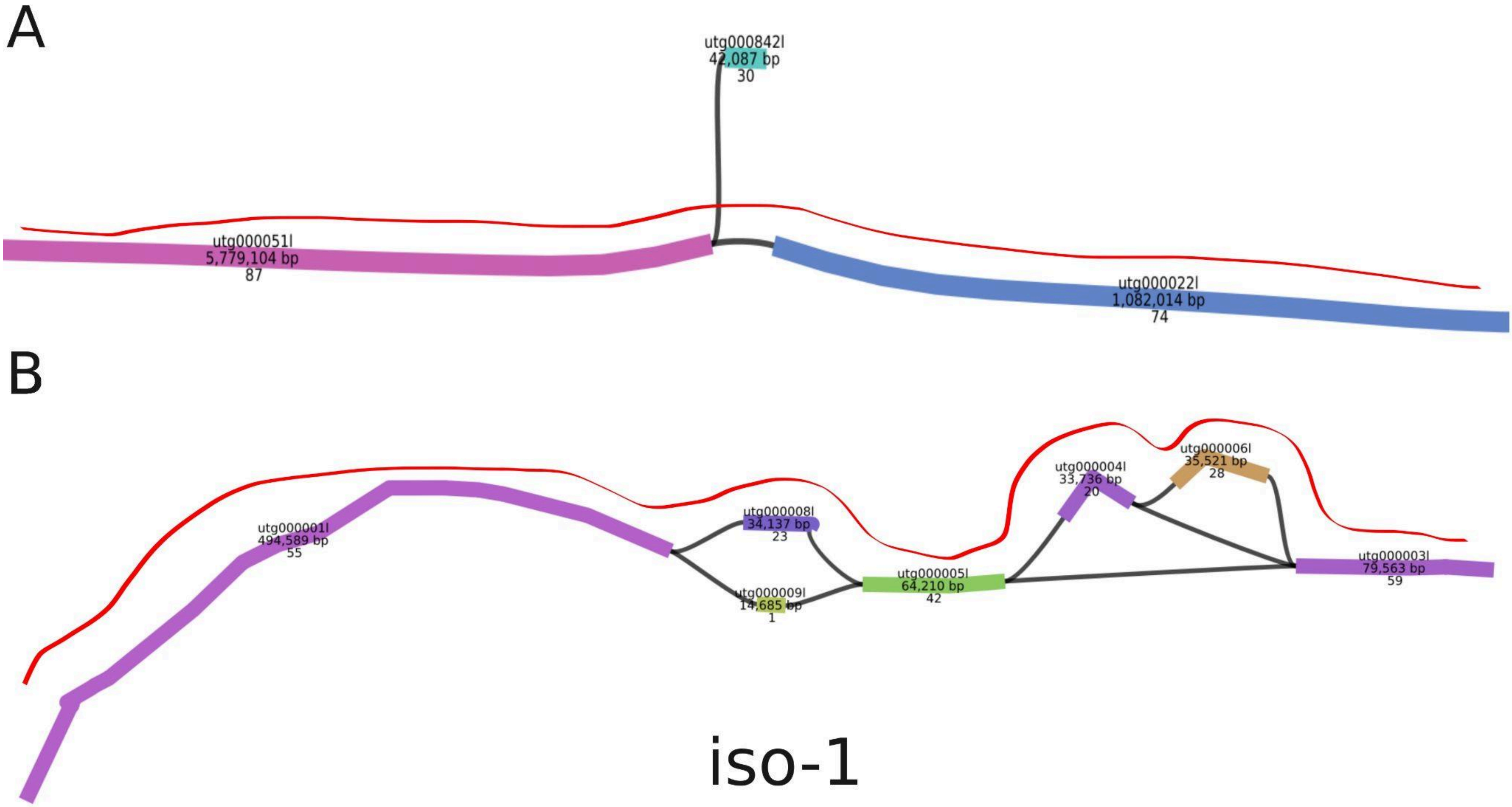


**Supplementary Fig 9A.** The 3 *unitigs* that contain the histone locus in the original iso-1 assembly. The 1st line of the node label is nodename. The 2nd line is the length of the node. The 3rd line is the depth (scales to number of reads covering that particular node). The red line indicates the possible path taken by the assembler to get through the locus.
**Supplementary Fig 9B.** The unitig graph of the targeted histone assembly for iso-1.The red line indicates the possible path taken by assembler to get through the locus.

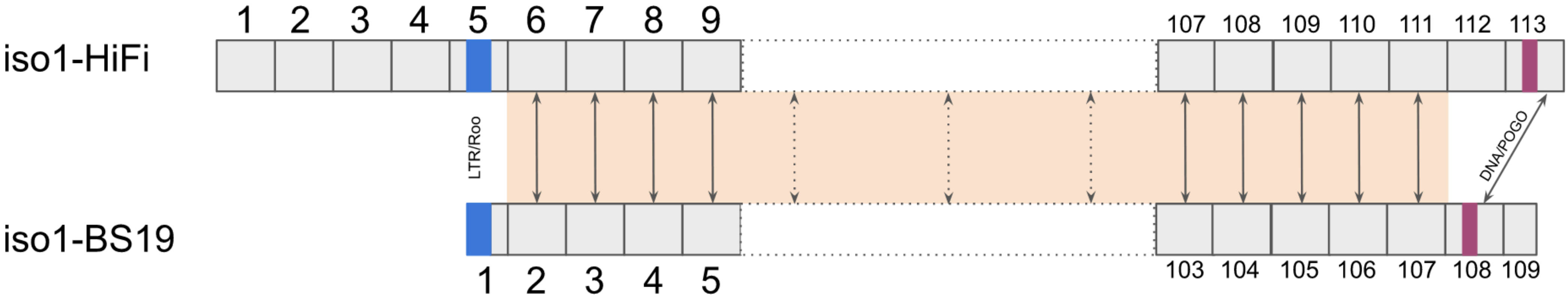


**Supplementary Fig 10.** A schematic representation of comparison between iso1-HiFi histone cluster vs iso-1 BS19 histone cluster.

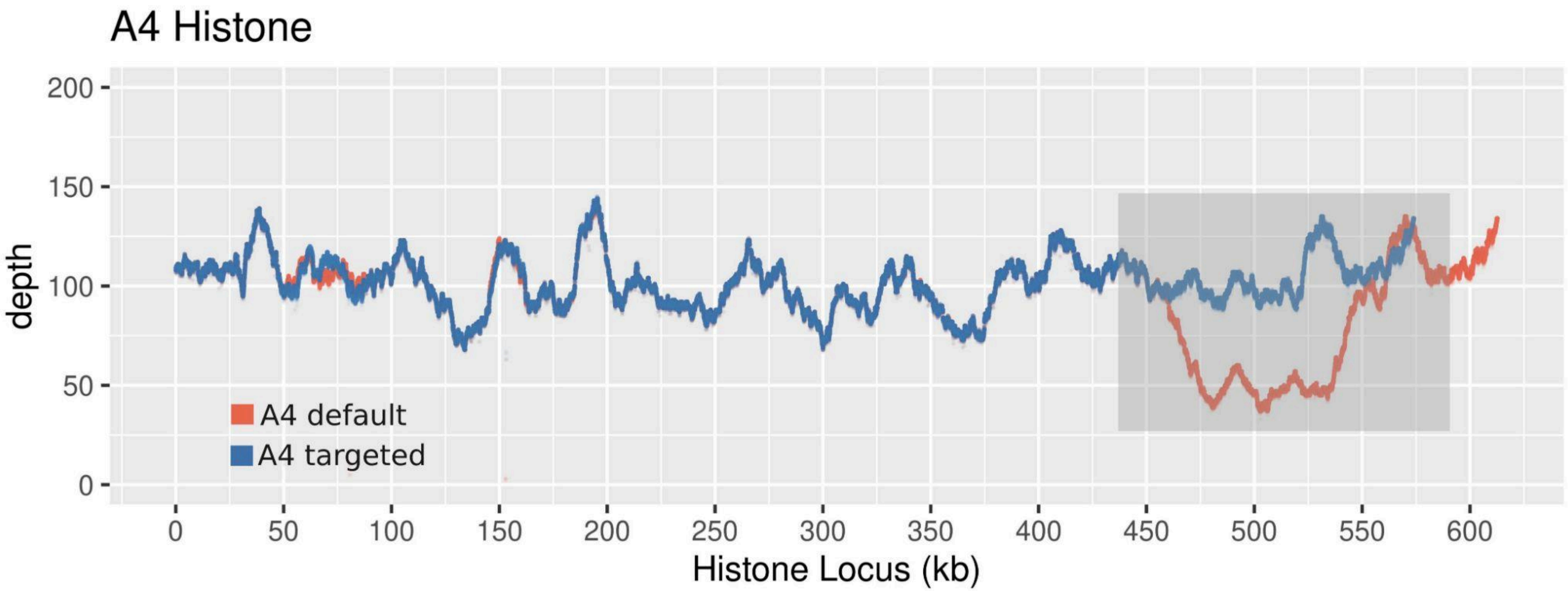


**Supplementary Fig 11.** The depth plots for A4 default and A4 targeted histone locus. There is a drop in coverage (~half) starting at ~425 kb to ~550 kb (gray box) in the default assembly. This tells us that a region present once in the genome is represented twice in the assembly. The depth plot for targeted assembly is relatively uniform, indicating no obvious/glaring mis-assemblies.

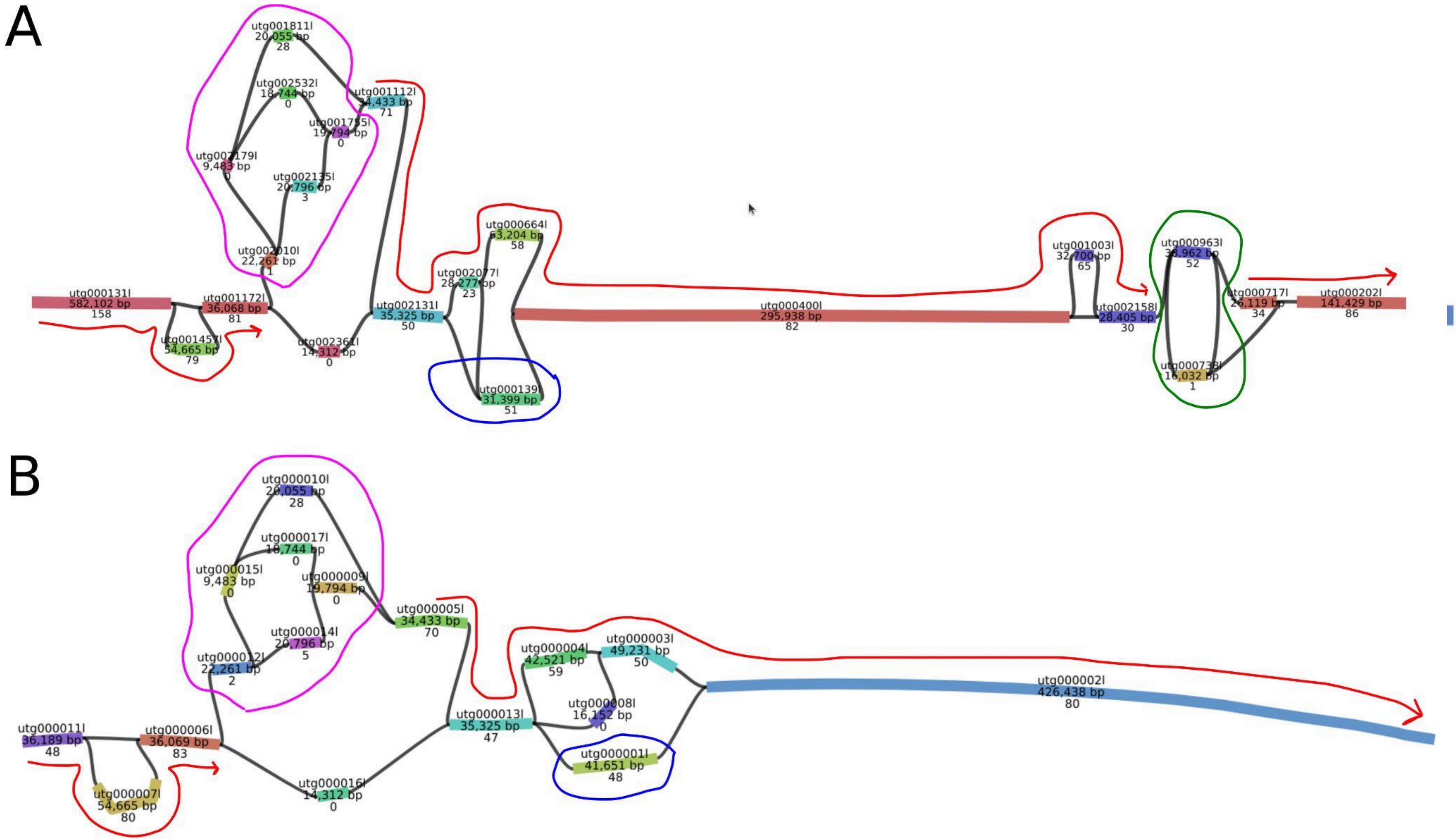


**Supplementary Fig 12 A & B.** The unitig graph for default (A) and targeted histone (B) assembly in A4. The 1st line of the node label is nodename. The 2nd line is the length of the node. The 3rd line is the depth. The green ~circle in A highlights the possible reason for drop in coverage seen in the depth plot (by visiting the nodes twice). The pink ~circle present in both graphs might represent a low coverage or complex region or segregating alleles . The red line indicates the possible path taken by the assembler to get through the locus. In both graphs, the high depth node highlighted by the blue circle doesn't get included in the primary contig. This might represent a high frequency segregating allele in the sequenced pool (since we sequenced ~200 diploid males).

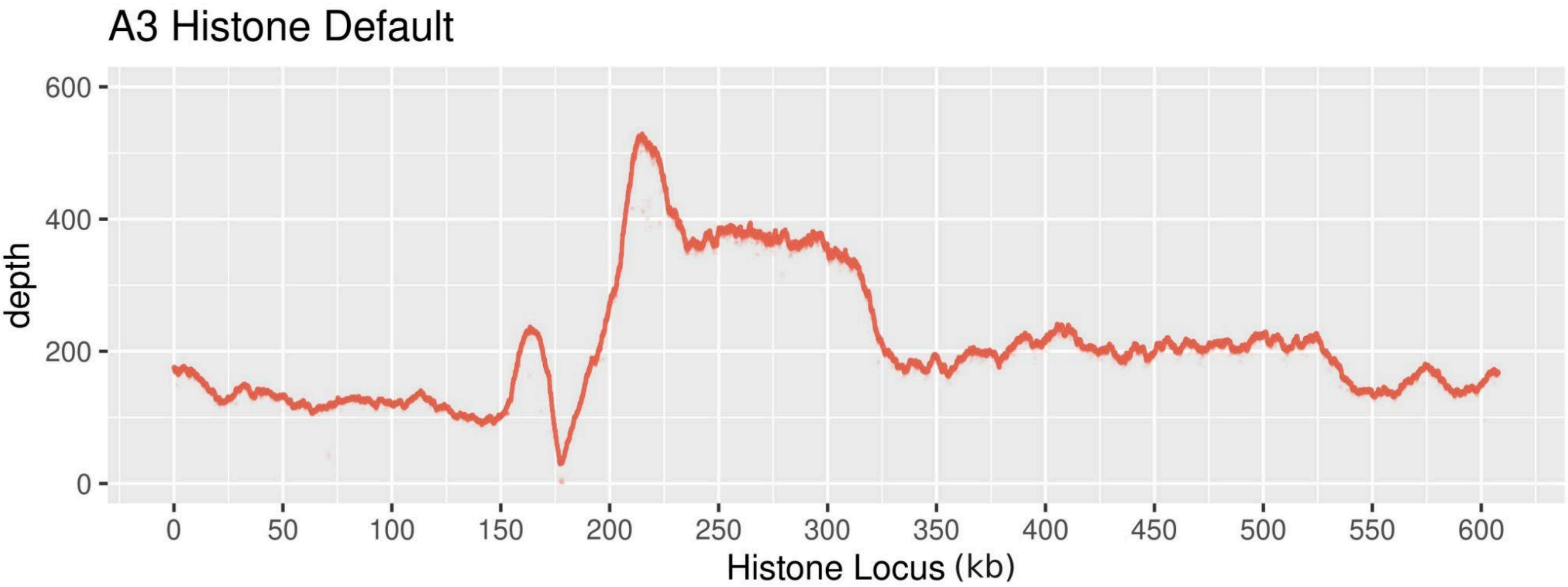


**Supplementary Fig 13.** The depth plot for A3 default histone locus. There are multiple issues with the assembly starting at ~150 kb.

A

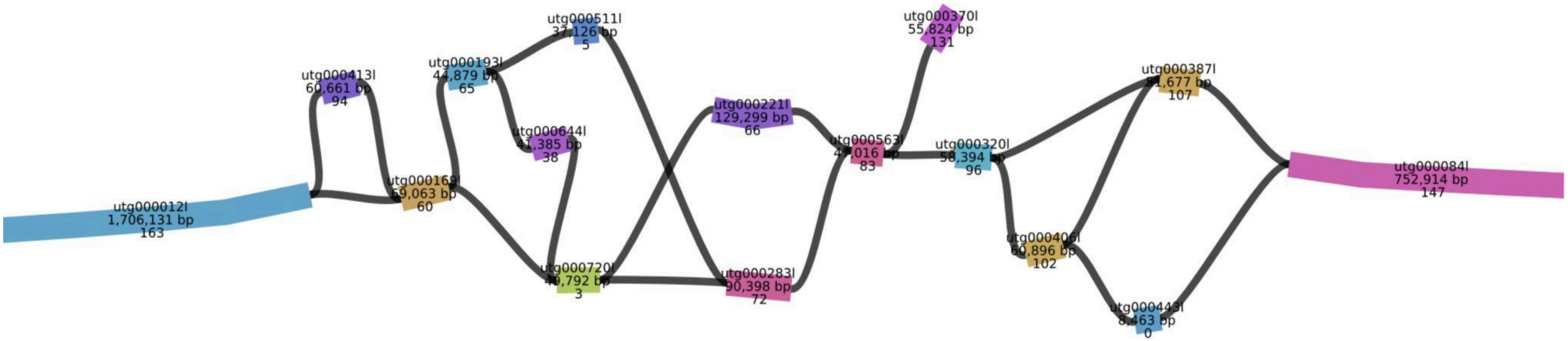


B

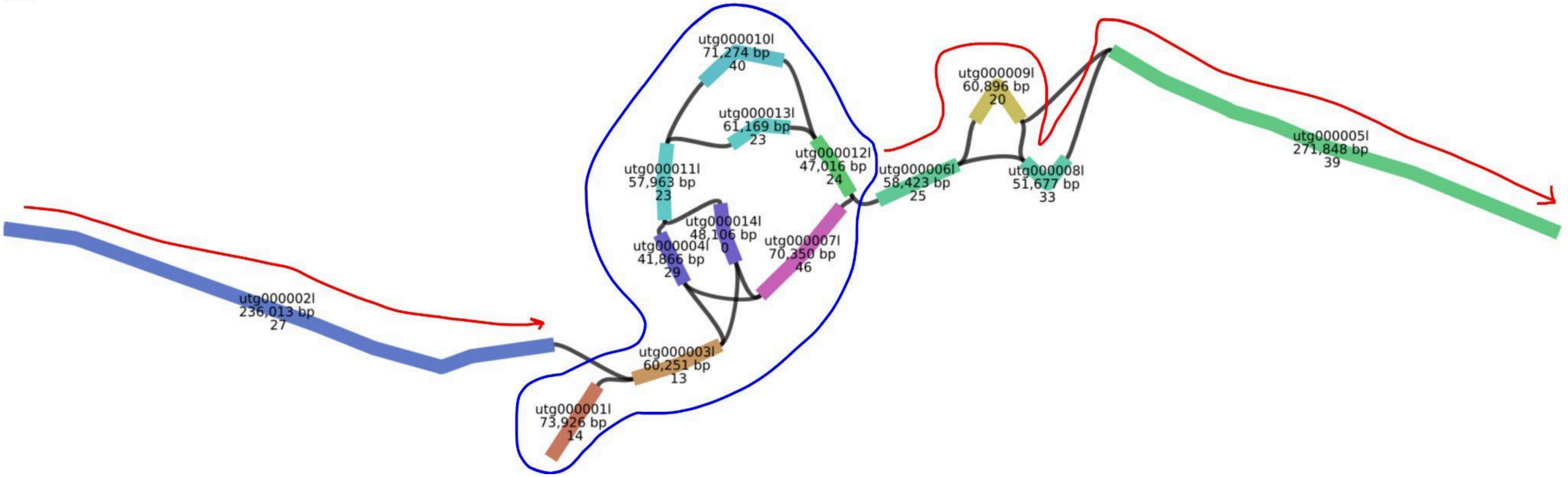


**Supplementary Fig 14 A & B.** The unitig graph for default (A) and longest-40X targeted (B) assemblies. The red line in B represents the well resolved parts of the graph. The blue circle represents the complex region in the middle (path through it cannot be cannot be unambiguously determined).

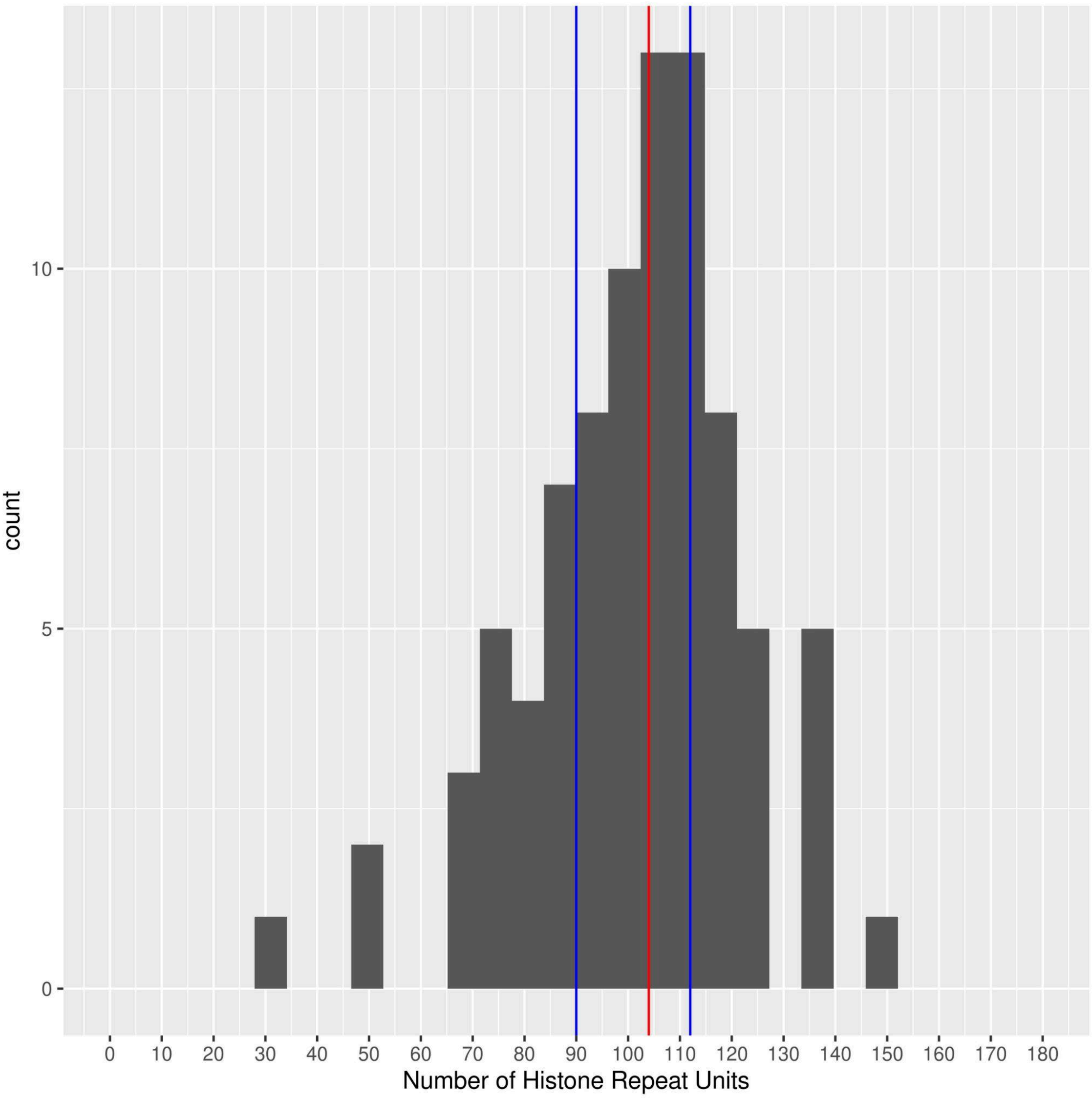


**Supplementary Fig 15.** Histone copy number distribution of the GDL strains. The red line indicates the median and the blue lines represent the first (Q1) and the third (Q3) quartile.

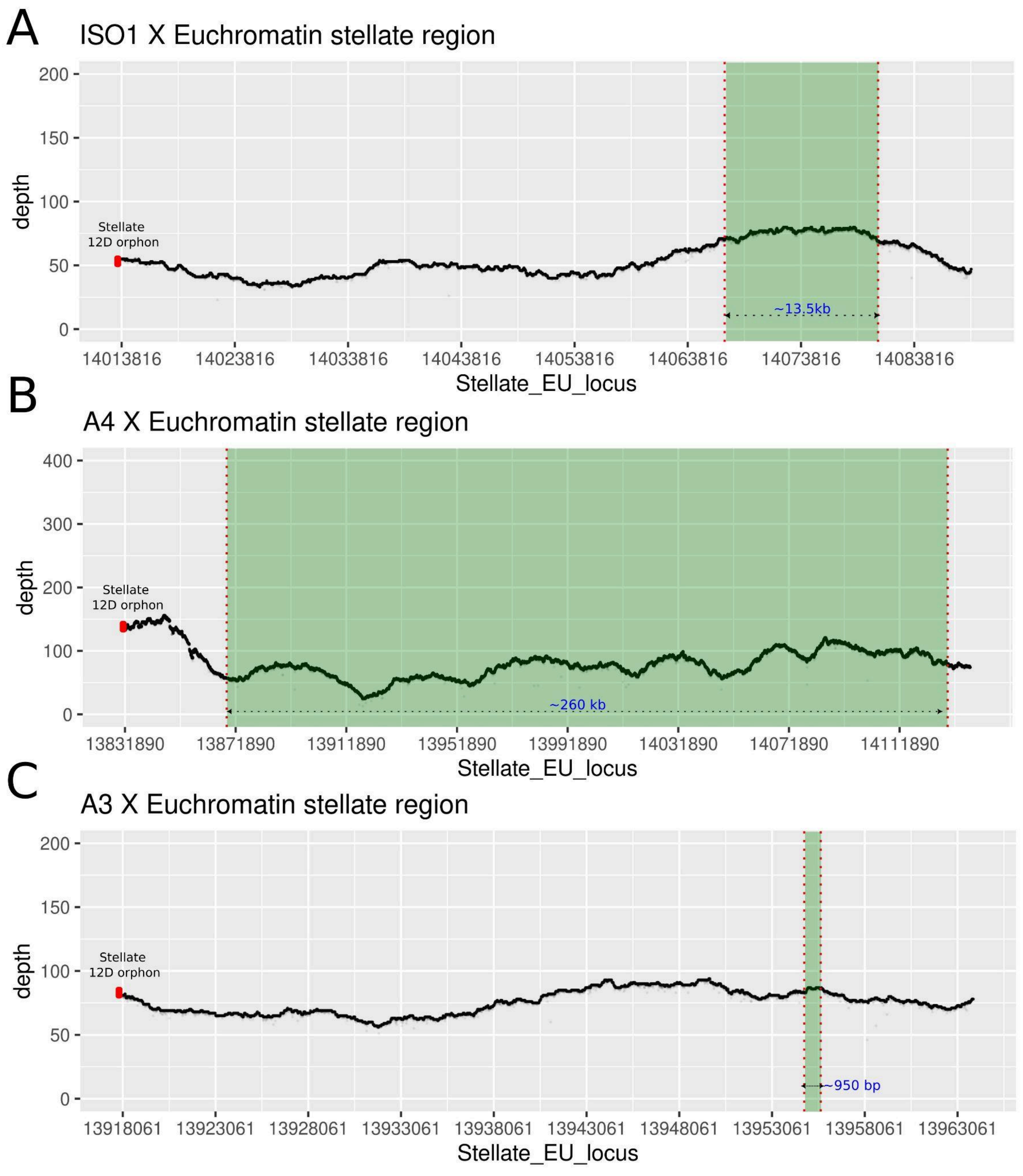


**Supplementary Fig 16 A B C.** HiFi depth plots for iso-1 (A), A4 (B) and A3 (C) for the euchromatin stellate locus. The red mark indicates the start of *Stellate 12D orphan copy*. The green rectangle highlights the stellate proper tandem cluster.

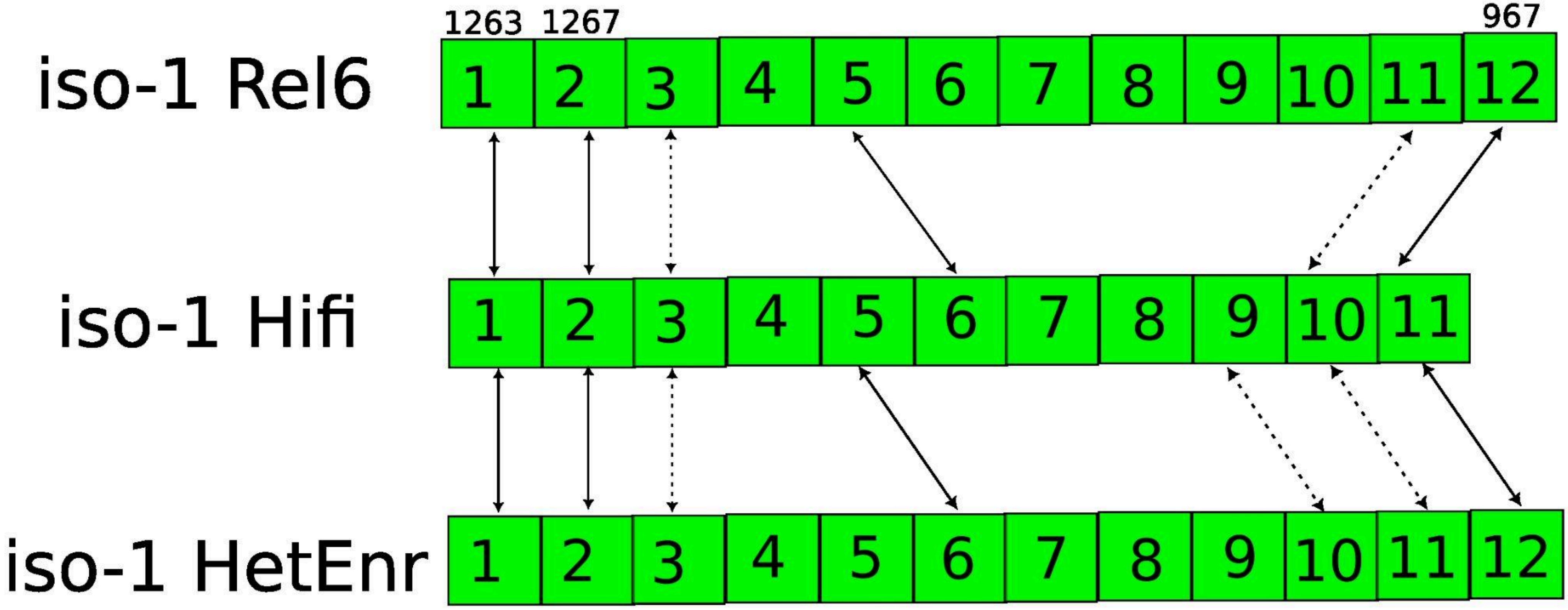


**Supplementary Fig 17.** A schematic representation of euchromatic stellate clusters in the 3 iso-1 assemblies and their comparison. The solid arrows represent the anchors identified using phylogenetic methods. The dashed arrow indicates possible/probable anchors marked manually using proximity (synteny) information to the phylogenetically identified anchors (solid arrows). The numbers on the top represent the lengths of the stellate units (which deviate from the canonical 1269 bp unit).

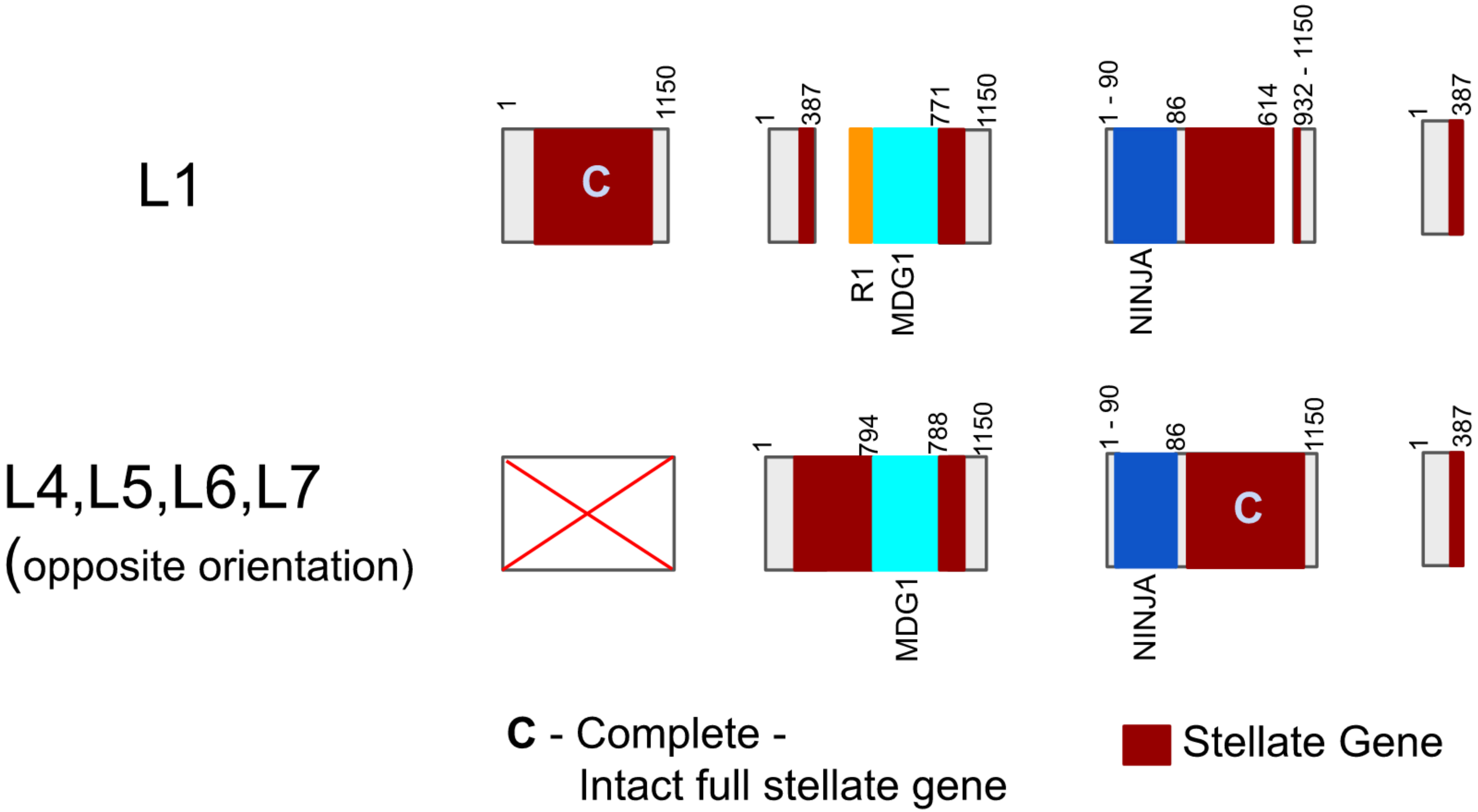


**Supplementary Fig 18.** A schematic representation of Type_1 locus in iso-1. The numbers on the top are coordinates corresponding to the full length (1150) repeat unit for each copy. R1, NDG1 and NINJA represent TE insertions. The red box marks the *stellate gene* embedded within the whole 1150 repeat unit. **C** indicates a full length *Stellate* repeat unit/gene.

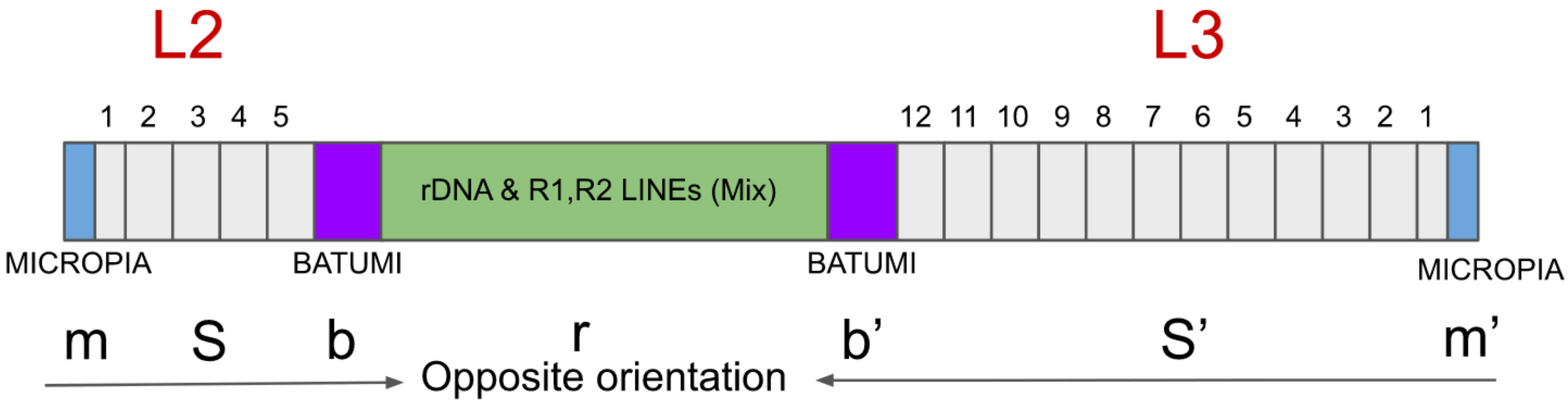


**Supplementary Fig 19.** A schematic representation of Type_2 loci (L2 and L3) and their placement with respect to each other. The numbers on the top correspond to a particular copy in the tandem array. MICROPIA and BATUMI are LTR TEs. m,s,b,r are shorthand used to describe m-MICROPIA, S-Stellate array, b-BATUMI and r-rDNA & R1,R2 LINEs (Mix). ‘ indicates opposite orientation. The Unit labeled as 1 has a partial deletion (highlighted by short box length).

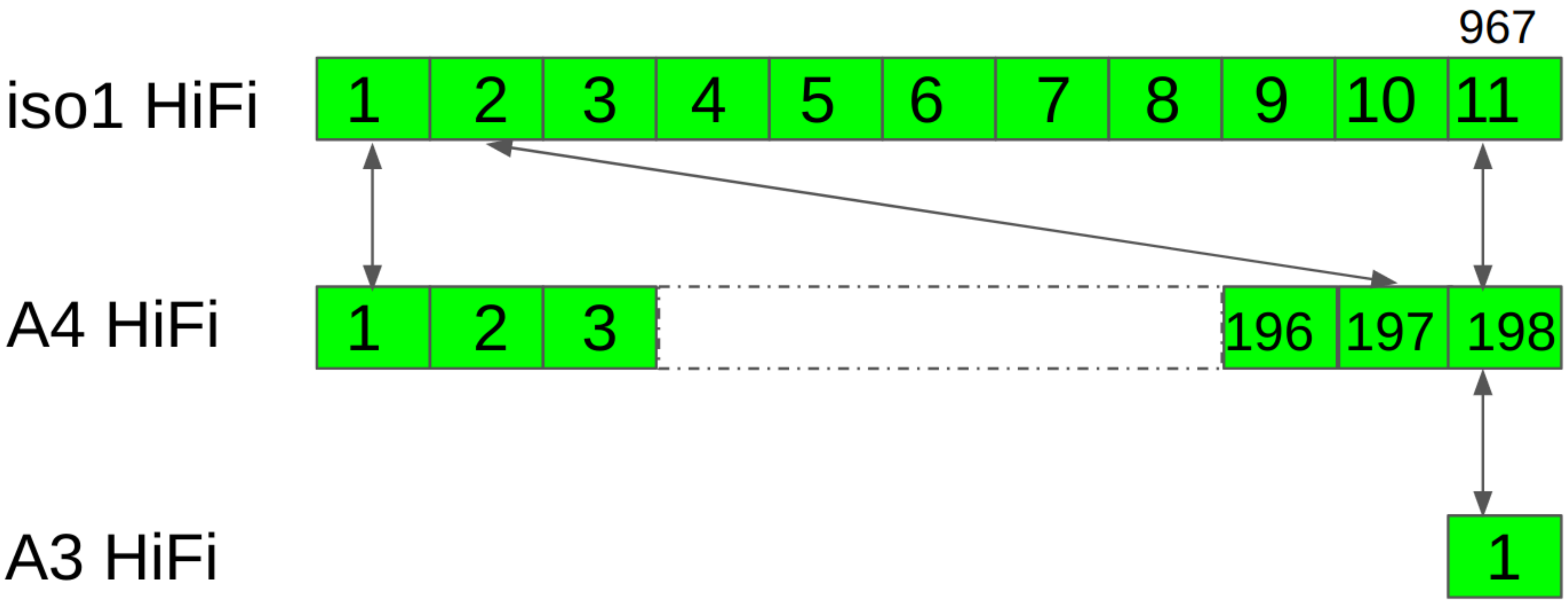


**Supplementary Fig 20.** A schematic representation of euchromatic stellate clusters in iso-1,A4 and A3 assemblies and their comparison. The numbers in the box designate the unit in an array. The dotted gray box in A4 represents unit 4-195. The solid arrows indicate anchors identified using phylogenetic approaches. The number (967) on top of last unit represents the most proximal unit (with 302 bp deletion)

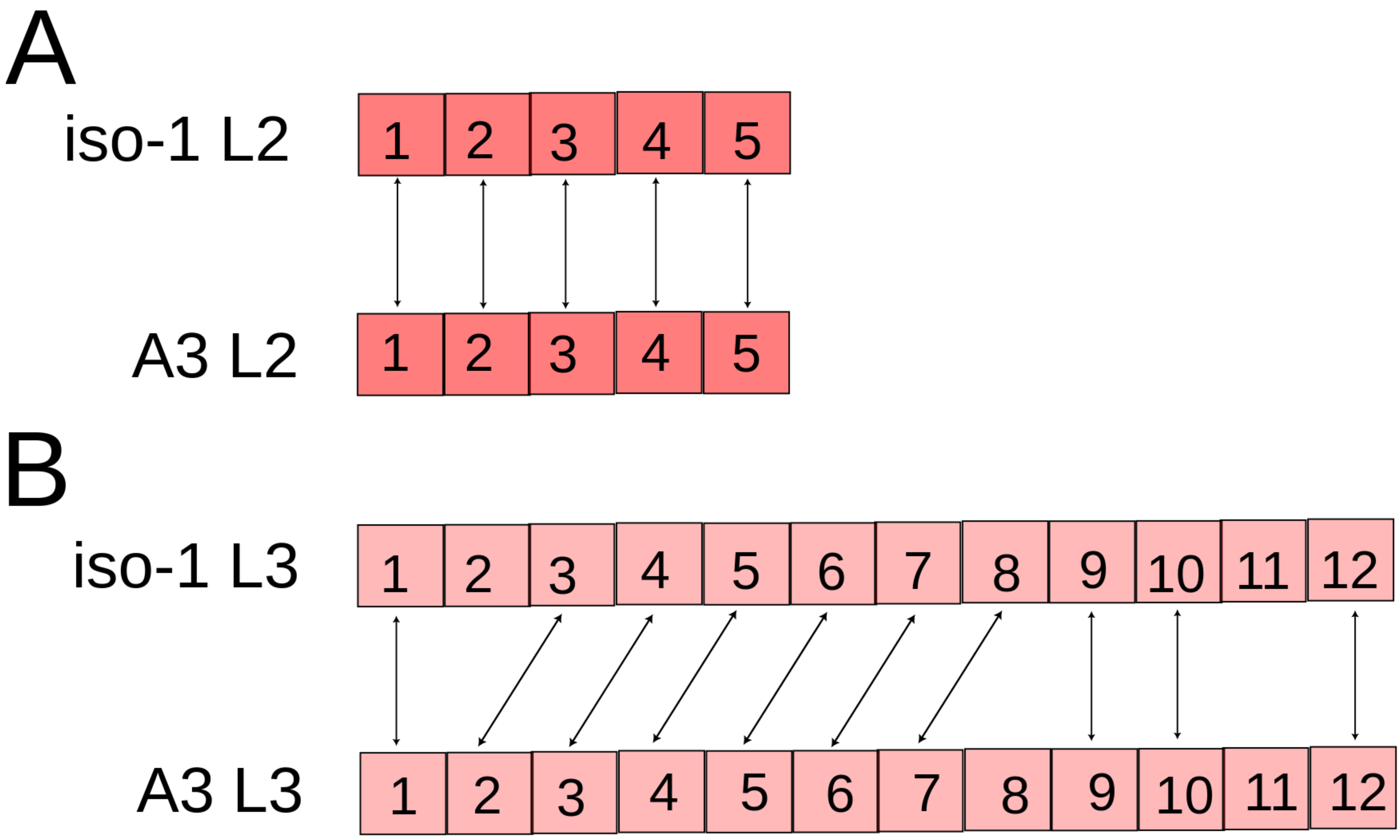


**Supplementary Fig 21.** A schematic representation of heterochromatic stellate clusters in iso-1 and A3 assemblies and their comparison. There is one to one correspondence for all units in L2 **(A)**. iso-1 L3 and A3 L3 are mostly similar with some minor differences **(B)**.

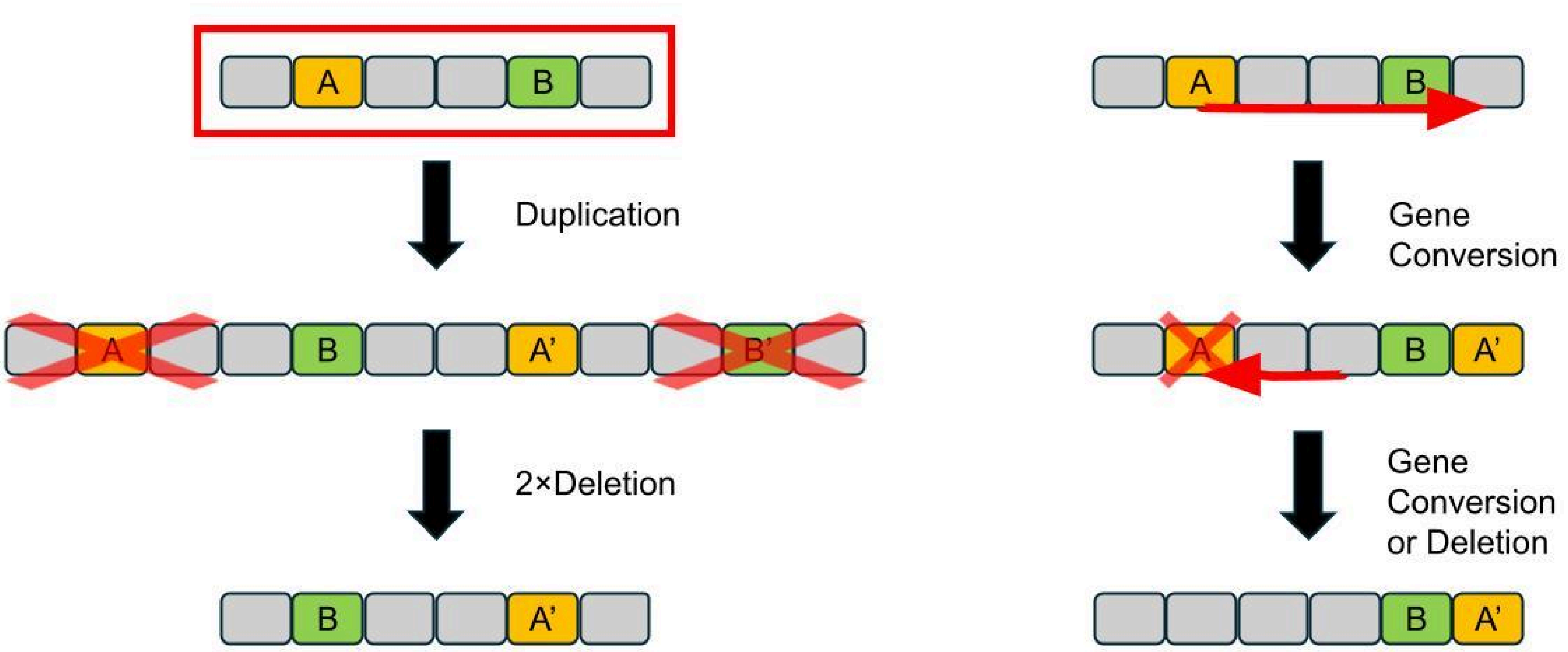


**Supplementary Fig 22**. Two possible scenarios demonstrating mutational steps that can result in anchor points swapping their relative order in the array.

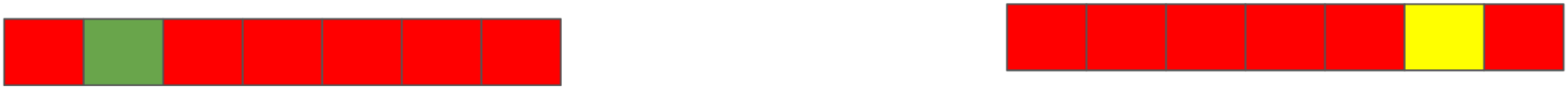

Suppose two new variants (green and yellow) arise on two separate haplotypes in a population. After some time has passed, two haplotypes are again sampled from the population. You either see **A** or **B**

A

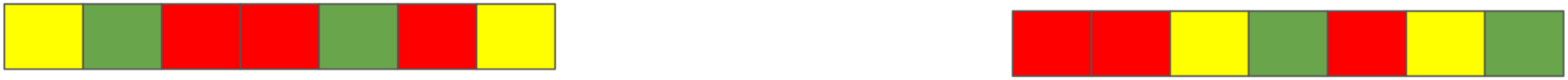

If there is significant recombination between different haplotypes (crossing over or gene conversion) the duplications will be mostly spread within the haplotype and present in somewhat equal proportions in all haplotypes

B

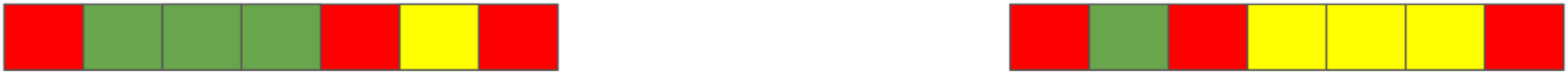

But we see something like this .. the duplicates are mostly together and they are present in different proportions in different haplotypes. This means the rate of intrachromosomal exchange (recombination within the same haplotypic lineage) is more than interchromosomal exchanges. If the rate of interchromosomal recombination (specifically crossing over here) was similar to intrachromosomal one it would quickly start unlinking the duplicates generated by intrachromosomal mechanisms.

**Supplementary Fig 23**. Simplified diagrams demonstrating two possible outcomes of spread of newly arisen variants in a population, given some rate of inter- and intra-chromosomal exchanges. Our observations (B) suggest that the rate of intrachromosomal recombination is significantly higher than interchromosomal one.

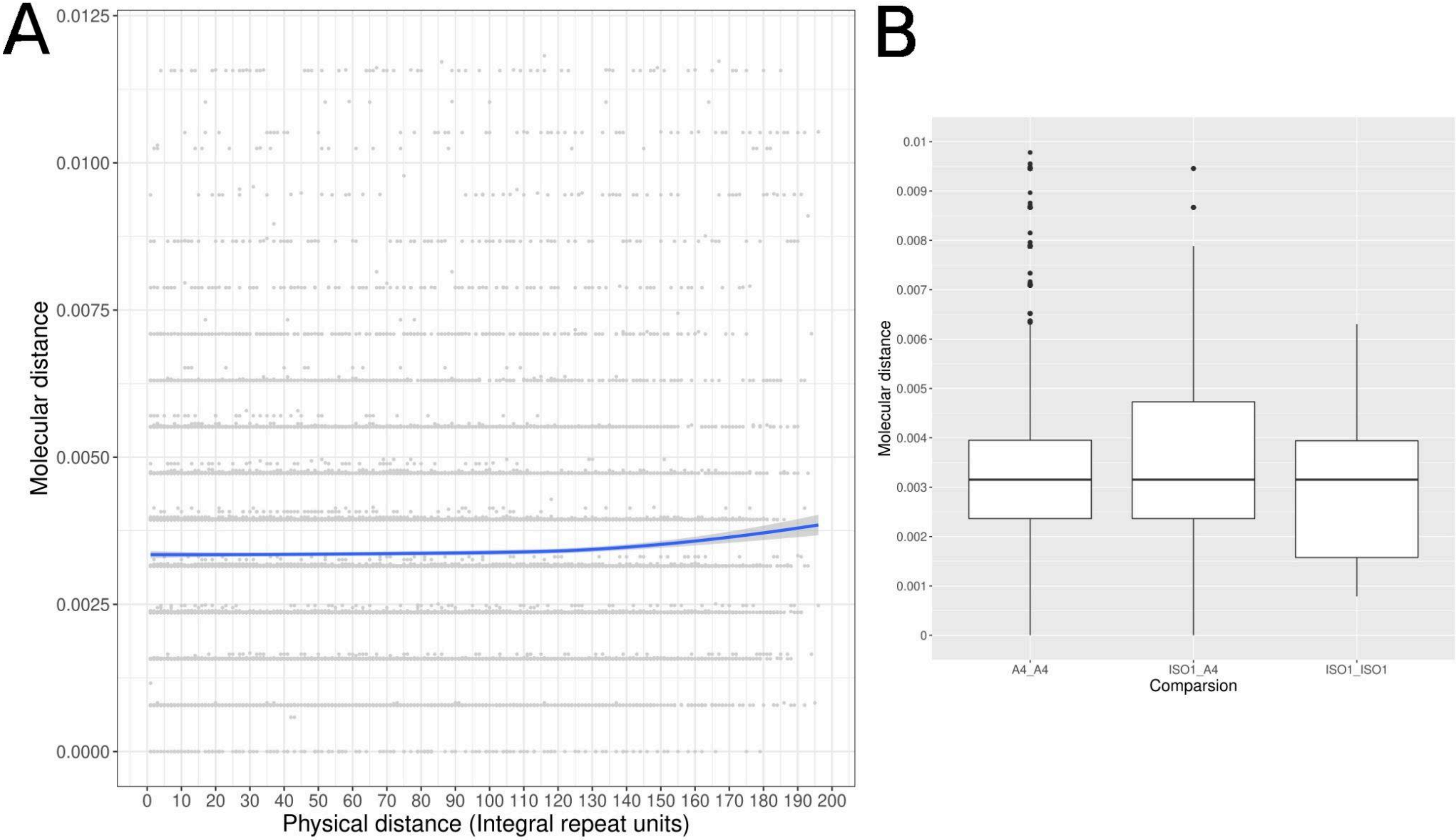


**Supplementary Fig 24. (A)** Physical vs molecular distance plot for A4 euchromatic stellate array. (Similar to Fig 4C) **(B)** Box plot comparing molecular distance between all pairwise units for within versus between array comparison for iso-1 and A4.
